# Maternal antibodies facilitate Amyloid-β clearance by activating Fc-receptor-Syk-mediated phagocytosis

**DOI:** 10.1101/2020.09.22.296376

**Authors:** Tomer Illouz, Raneen Nicola, Linoy Ben-Shuhan, Ravit Madar, Arya Biragyn, Eitan Okun

## Abstract

Down Syndrome (DS) features a life-long overexpression of the *APP* and *DYRK1A* genes, leading to a cognitive decline mediated by Amyloid-β (Aβ) overproduction and tau hyper-phosphorylation. As DS can be diagnosed in utero, maternally transferred anti-Aβ antibodies might promote removal of early accumulation of Aβ from the CNS. A DNA-vaccine expressing Aβ_1-11_ was delivered to wild-type female mice, followed by mating with 5xFAD males, which exhibit early Aβ plaque formation, similar to individuals with DS. Maternal Aβ-specific antibodies provided transgenic offspring with passive immunization against Aβ via the placental and subsequently lactation. Maternal antibodies reduced cortical Aβ levels 4 months after antibodies were undetectable, along with alleviating short-term memory deficits and activation of the FcγR1/Syk/Cofilin pathway in microglia. Sera from immunized dams facilitated Aβ clearance by microglia in a Syk-dependent manner. These data suggest that maternal anti-Aβ immunization is a potential strategy to alleviate cognitive decline in individuals with DS.

## Introduction

Down syndrome (DS), caused by trisomy of chromosome 21, is the most prevalent genetic cause for intellectual disability, affecting ∼1 in every 750 live births (Presson et al., 2013). The life expectancy of individuals with DS has risen significantly in the past several decades due to improved medical treatment of various comorbidities. Nevertheless, individuals with DS have a shortened life-expectancy and accelerated aging compared to the general population (Carfi et al., 2014). DS shares common pathological symptoms with Alzheimer’s disease (AD) (Choong, Tosh, Pulford, & Fisher, 2015), as individuals with DS experience AD-related cognitive decline in their 4^th^ and 5^th^ decades of their lives (Choong et al., 2015; Wiseman et al., 2015). This is preceded by a significant and widespread AD-related pathology, detected as early as at the age of 12 years (Burger & Vogel, 1973), which includes intraneural neurofibrillary tangles, amyloid-β (Aβ) angiopathy, extracellular Aβ neuritic plaques (Alhajraf, Ness, Hye, & Strydom, 2019), and microgliosis (Heneka, 2019; Raha-Chowdhury et al., 2018; Tejera & Heneka, 2019). This is in part due to over-expression of the *APP* and *DYRK1A* genes, located on the triplicated chromosome 21 (Korbel et al., 2009), although Aβ deposition is also promoted by non-*APP* mediated mechanisms in this pathology (Wiseman et al., 2015; Wiseman et al., 2018). In this respect, Aβ-related pathology in DS shows high similarity to early-onset AD (EOAD), which results from an over-expression or mutation in the genes for presenilin-1, 2, or amyloid precursor protein (APP) (Chan, Furukawa, & Mattson, 2002). Thus, Aβ pathology is etiologically and mechanistically shared between EOAD and DS (Barone, Head, Butterfield, & Perluigi, 2017; Butterfield & Perluigi, 2018; Tramutola et al., 2018). As DS is typically diagnosed towards the end of the first trimester of the pregnancy, individuals with DS present a unique group that can benefit from an early intervention directed at Aβ pathology.

Maternal vaccination can be administered during pregnancy against influenza infection, Tetanus toxoid, reduced diphtheria toxoid, and acellular pertussis. Pneumococcal, Meningococcal, and hepatitis A and B vaccines may be given in pregnancy in specific populations. All the above-mentioned vaccinations can also be administered post-partum, during breastfeeding, or both (ACOG, 2018; Munoz & Jamieson, 2019). Thus, in many countries, maternal vaccination is a routine procedure during pregnancy, post-partum, and during breastfeeding.

AβCoreS is a DNA vaccine coding Aβ_1-11_ (a B-cell epitope) fused to a Hepatitis-B surface antigen (HBsAg) and a Hepatitis-B core antigen (HBcAg), both contain multiple T-helper epitopes that help facilitate antibody (Ab) production (Olkhanud et al., 2012). The AβCoreS construct was previously shown to delay cognitive decline and reduce human Aβ pathology in the 3xtgAD mouse model of AD (Olkhanud et al., 2012). A modified vaccine targeting murine Aβ, which is triplicated in the Ts65Dn mouse model of DS (Gupta, Dhanasekaran, & Gardiner, 2016), was also shown to induce anti-Aβ Ab production that facilitates clearance of soluble oligomers and small extracellular inclusions of Aβ from the hippocampus and cortex of Ts65Dn mice (Illouz, Madar, Biragyn, & Okun, 2019). This was correlated with reduced neurodegeneration and restoration of the homeostatic phenotype of microglia and astrocytes (Illouz et al., 2019). These findings support the notion that immunotherapy against Aβ can slow the progression of dementia in DS, and that an anti-murine Aβ vaccination is safe to use in mice, as vaccinated wild-type (WT) mice exhibited no cognitive or other behavioral abnormalities that could indicate neurotoxicity (Illouz et al., 2019).

In the current study, we hypothesized that maternal vaccination at the developmental and post-natal stages coupled with an active vaccination post-partum would yield a continuous immune coverage against Aβ and a potent effect on AD-related neuropathology and dementia in DS.

Mouse models of trisomy-21 successfully recapitulate several aspects of DS, such as cholinergic neurodegeneration, elevated levels of Aβ and phospho-tau, and impaired cognitive ability (Rueda, Florez, & Martinez-Cue, 2012). However, these current DS models critically fail to recapitulate extra-cellular Aβ plaque accumulation (Illouz et al., 2019; Wiseman et al., 2018). Indeed, several DS models, including the Ts65Dn strain, overexpress murine Aβ, which does not aggregate as human Aβ, potentially due to three amino-acids differences in the N-terminus. These differences have been reported to modulate the binding of metal ions, which can influence the fibrillogenesis of Aβ peptides (Atwood et al., 1998). The replacement of His-13 for Arg in rodent Aβ disrupts a metal coordination site, rendering the rodent peptide less prone to zinc-induced aggregation *in vitro*. Moreover, this region is essential for the specificity of amyloid interactions (Jankowsky et al., 2007). Thus, mouse models that encompass triplication of murine APP exhibit elevated levels of APP protein and soluble Aβ but not plaque pathology. Humanized models of DS such as the Tc1 strain also lack plaque pathology due to a nonfunctional human *APP* gene (Gribble et al., 2013). Accordingly, these models lack plaque-associated microglial pathology (Illouz et al., 2019; Keren-Shaul et al., 2017; Krasemann et al., 2017). These considerations led us to use the 5xFAD strain that encompasses five mutations associated with Aβ pathology in EOAD (Oakley et al., 2006). Indeed, the 5xFAD model has low construct validity in modeling DS, nonetheless, its face validity in recapitulating early Aβ plaque pathology in DS is satisfactory compared with other AD and DS mouse models.

Herein, WT female mice were vaccinated against hAβ_1-11_ or sham at 8w of age (Fig. 1A). Vaccinated mice were then crossed with male 5xFAD mice to produce maternally vaccinated transgenic and WT offspring. Following weaning, transgenic offspring were actively vaccinated against Aβ_1-11_ or sham. The offspring were then tested for cognitive capacity at 4m and were sacrificed for neuropathology assessment at the age of 5m (Fig. 1A). Aβ-specific antibodies from vaccinated mice directly affected the phagocytosis capacity of primary microglial cells in a Syk-dependent manner. Our findings provide evidence for a long-term effect of maternal vaccination on Aβ-related pathology, long after maternal Abs were not detectable in the circulation of the offspring.

**Fig. 1.**
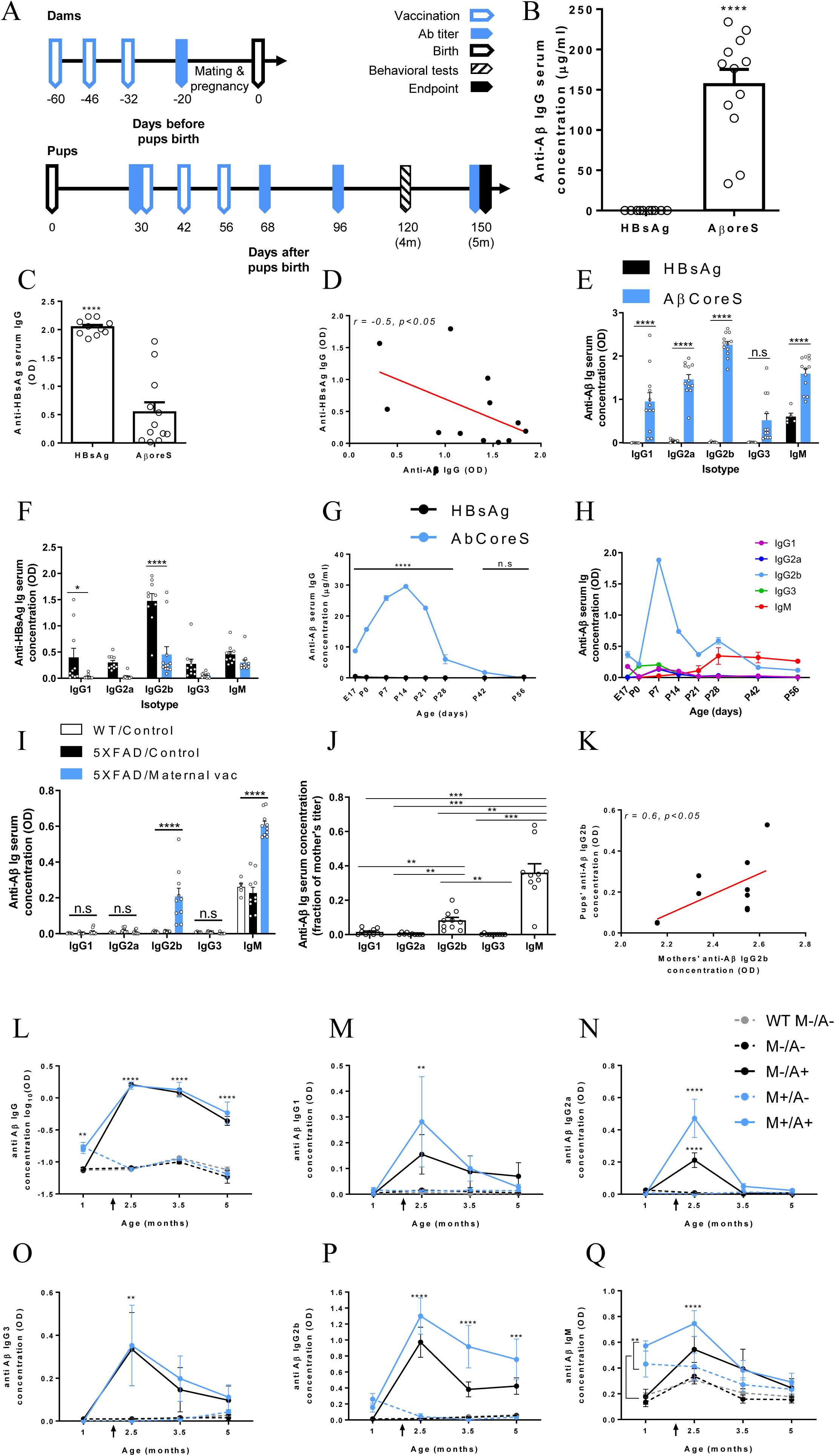
Maternally induced anti-Aβ antibodies cross the placenta and lactation into the circulation of 5xFAD fetuses and newborns. (A) Study design for dams (upper timeline) and pups (bottom timeline). Serum anti-Aβ antibodies were measured in vaccinated dams (B-F), offspring features, and newborns (G-K), and following active vaccination in the offspring (L-Q). (B) Anti-Aβ IgG titer in AβCoreS and HBsAg vaccinated dams. (C) Anti-HBsAg IgG titer in AβCoreS and HBsAg vaccinated dams. (D) A negative correlation between anti-Aβ IgG and anti-HBsAg IgG in vaccinated dams (E) Anti-Aβ Ig isotypes in AβCoreS and HBsAg vaccinated dams. (F) Anti-HBsAg Ig isotypes in AβCoreS and HBsAg vaccinated dams. (G) Maternally induced anti-Aβ Abs cross the placenta and lactation to the circulation of fetuses and newborns. (H) IgG2b is the most transferable isotype via the placenta and lactation. (I) Antibody titer was measured in the serum of maternally vaccinated and control 1m-old mice. (J) Ig isotypes in the serum of 1m-old offspring, normalized to mothers’ titer before pregnancy. (K) Offspring IgG2b is positively correlated with maternal IgG2b prior to pregnancy. Maternally and sham vaccinated mice were actively vaccinated at 1m of age against Aβ_1-11_. Antibody titer was measured at 2.5, 3.5, and 5m. (L) log_10_OD of total IgG concentration in the serum. Serum levels of (M) IgG1, (N) IgG3, (O) IgG2a, (P) IgG2b, and (Q) IgM. *P<0.05, **P<0.01, ***P<0.001, ****P<0.0001, unpaired t-test, linear regression, Pearson’s correlation, one-way ANOVA, two-way ANOVA, repeated measures two-way ANOVA.

## Results

### Maternal anti-Aβ antibodies are transferred to 5xFAD fetuses and newborns

Female C57BL/6j mice (8w-old) were immunized against Aβ_1-11_ using the AβCoreS-DNA construct or HBsAg as control (Fig. 1A). Vaccinated females produced a high anti-Aβ Ab titer compared with controls (156.4±18.98, 0±0μg/ml, respectively, P<0.0001, Fig. 1B). Both mouse groups generated anti-HBsAg titers, however, mice vaccinated with the HBsAg alone produced a higher titer than mice vaccinated with AβCoreS that includes Aβ_1-11_ and HBsAg domains (2.04±0.04, 0.56±0.17 OD, respectively, P<0.0001, Fig. 1C), possibly due to epitope competition. This is further indicated by a negative correlation of *r*=-0.5 found between anti-Aβ and -HBsAg Ab levels (P<0.05, Fig. 1D). The AβCoreS vaccine elicited high IgG1, IgG2a, IgG2b, and IgM titers against Aβ (P<0.0001, Fig. 1E). HBsAg alone induced higher IgG1 levels (0.39±0.17, 0.02±0.01 OD, respectively, P<0.05, Fig. 1F) and higher IgG2b levels (1.47±0.13, 0.45±0.15 OD, respectively, P<0.0001, Fig. 1F) compared with AβCoreS-vaccinated mice.

Following vaccination, WT females were mated with 5xFAD males to produce hemizygous transgenic and WT offspring. For Abs analysis, circulating blood was taken from 5xFAD fetuses at E17, P0, P7, P14, P21, P28, P42, and P56. Maternal Anti-Aβ Abs crossed the placenta to the fetuses, as early as E17, compared with controls (8.73±0.31, 0.46±0.17μg/ml, respectively, P<0.0001 Fig. 1G). Strikingly, these Abs crossed to a greater extent via lactation of the colostrum (P0) and milk (P7-P21). Ab levels gradually increased in vaccinated fetuses compared with controls, with the progression of breastfeeding reaching a peak at P14 (29.57±0.65, 0±0μg/ml, respectively, P<0.0001, Fig. 1G). Following weaning at P28, Ab levels decreased to those of controls. At P42, no Ab difference was observed between the two groups (1.76±0.65, 0±0μg/ml, respectively, P=0.12, Fig. 1G). IgG2b was the most prevalent Ig isotype to transfer via the placenta and lactation, peaking at P7 (Fig. 1H).

### Active immunization of maternally vaccinated 5xFAD pups induces mostly anti-Aβ IgG2b Abs

We next assessed whether maternal vaccination is comparable to active vaccination at a young age. Prior to active vaccination, elevated levels of Aβ-specific IgG2b and IgM were found in maternally vaccinated pups compared with controls (P<0.0001, Fig. 1I). Normalizing the pups’ isotype concentration to those of their dams revealed that IgG2b levels were 8.3% of the mothers’ circulating concentration, significantly higher than IgG1, IgG2a, and IgG3 (P<0.01, Fig. 1J). A significant positive correlation of *r*=0.6 was found between dams’ and pups’ IgG2b levels (P<0.05, Fig. 1K). Normalized IgM levels were also high in pup circulation reaching 36% of the mothers’ concentration, higher than all other isotypes (P<0.01, Fig. 1J).

At 1m of age, both groups of Aβ_1-11_ and sham maternally vaccinated pups received either Aβ_1-11_ or sham active vaccination, to create four experimental groups: No treatment M-/A-, maternal vaccination only M+/A-, active vaccination only M-/A+, and combined maternal and active vaccination M+/A+. A control WT group was administered with the control vaccine only for behavioral and neuropathological baseline (Fig. 1A). Blood Ab titers at 1m reflected maternal IgG, whereas Ab titers at 2.5, 3.5, and 5m reflected humoral response to active vaccination. Following active vaccination, total IgG levels in the M-/A+ and M+/A+ groups dramatically increased compared with M-/A-, M+/A-, and WT groups (0.2±0.02, 0.19±0.05, -1.09±0.01, -1.12±0.02, -1.11±0.09 log_10_ OD, P<0.0001, respectively, Fig. 1L). The yield of active vaccination in the M+/A+ group was 18.24±4.09- fold the concentration of maternal Abs and remained high at 5m (P<0.0001, Fig. 1L). Exposure to maternal Abs in the M+/A- group was limited up to 1m of age, following which Ab levels were undetectable (Fig. 1L). In actively vaccinated offspring, IgG1, IgG2a, and IgG3 levels were higher than control at 2.5m (P<0.01, Fig. 1M, N, O, respectively) and decreased to baseline at later time-points. As expected, levels of IgG2b were elevated following vaccination and remained high until 5m of age in the M-/A+ and M+/A+ groups compared with the M-/A-, M+/A-, and WT groups (0.4±0.1, 0.75±0.25, 0.05±0.01, 0.03±0.005, 0.06±0.009 OD, respectively, P<0.001, Fig. 1P). Accordingly, IgM levels were elevated following vaccination among actively vaccinated mice at 2.5m (P<0.0001, Fig. 1Q), which decreased to baseline at later time-points. Summary of isotype levels is available in Supplementary, Fig. S1.

### Maternal and active vaccination rescue short-term memory abilities and normalizes exploratory behavior

To compare the effects of maternal and active vaccinations with that of Aβ on cognition, mice were tested in a battery of behavioral paradigms at 4m of age (Fig. 1A). Short-term memory capacity was assessed using the novel object recognition (NOR) test (Lueptow, 2017). While WT, M-/A+, and M+/A+ vaccinated 5xFAD mice exhibited a clear preference to the novel object throughout the experiment, unvaccinated mice showed no such preference after the first 20s of the trial (1.9±0.66, 2.45±0.31, 0.96±0.08, -4.4±0.03 cumulative discrimination index, respectively, P<0.0001, Fig. 2A_1-2_). M+/A- mice exhibited a preference to the novel object only during the first 60s (- 1.9±0.89, P<0.01 compared with M-/A-, P<0.0001 compared with M+/A+, Fig. 2A_1-2_), suggesting that maternal vaccination alone partially ameliorated deficits in short-term memory, while active vaccination and combined vaccination resulted in a full short-term memory restoration. M-/A- mice exhibited lower exploratory behavior in this task, as they spent less time near the familiar and novel objects together compared with M+/A- (0.39±0.12, 0.89±0.16, respectively, P<0.01, Fig. 2A_2_, B). No difference was observed in the spontaneous-alternation T-maze for short-term memory assessment (P=0.51, Fig. S2F). To rule out non-cognitive effects on short-term memory assessment, we also tested exploratory behavior. Similarly to WT controls, M+/A+ 5xFAD mice spent more time in the center of the open field (OF) arena (Seibenhener & Wooten, 2015), compared with M-/A-, M+/A-, and M-/A+ vaccinated 5xFAD mice (93.9±11.64, 74.88±12.2, 70.56±6.46, 65.34±9.87s, respectively, P<0.01, Fig. 2C_1-2_). Exploration speed and distance did not differ between groups (P=0.61, P=0.59, respectively, Fig. S2A-B). These results imply that a combination of maternal and active vaccination completely normalized exploratory behavior in 4m-old 5xFAD mice. To rule out anxiety-related confounds, mice were tested in the elevated zero-maze (EZM) (Shepherd, Grewal, Fletcher, Bill, & Dourish, 1994). The fraction of time spent in the open/close zones of the maze did not differ between groups (P=0.96, Fig. S2C), suggesting no effect on anxiety-related behavior. However, the number of entries to the anxiogenic open section of the EZM was lower among M-/A- mice compared with M+/A- (0.87±0.09, 1.33±0.08, respectively, Fig. 2D_1-2_). Exploration distance and speed did not differ between groups (P=0.15, P=0.15, respectively, Fig. S2D, E). These findings suggest that while no anxiety-like behavior is found, maternal and active vaccinations enhance exploratory behavior among 5xFAD mice in the EZM.

**Fig. 2.**
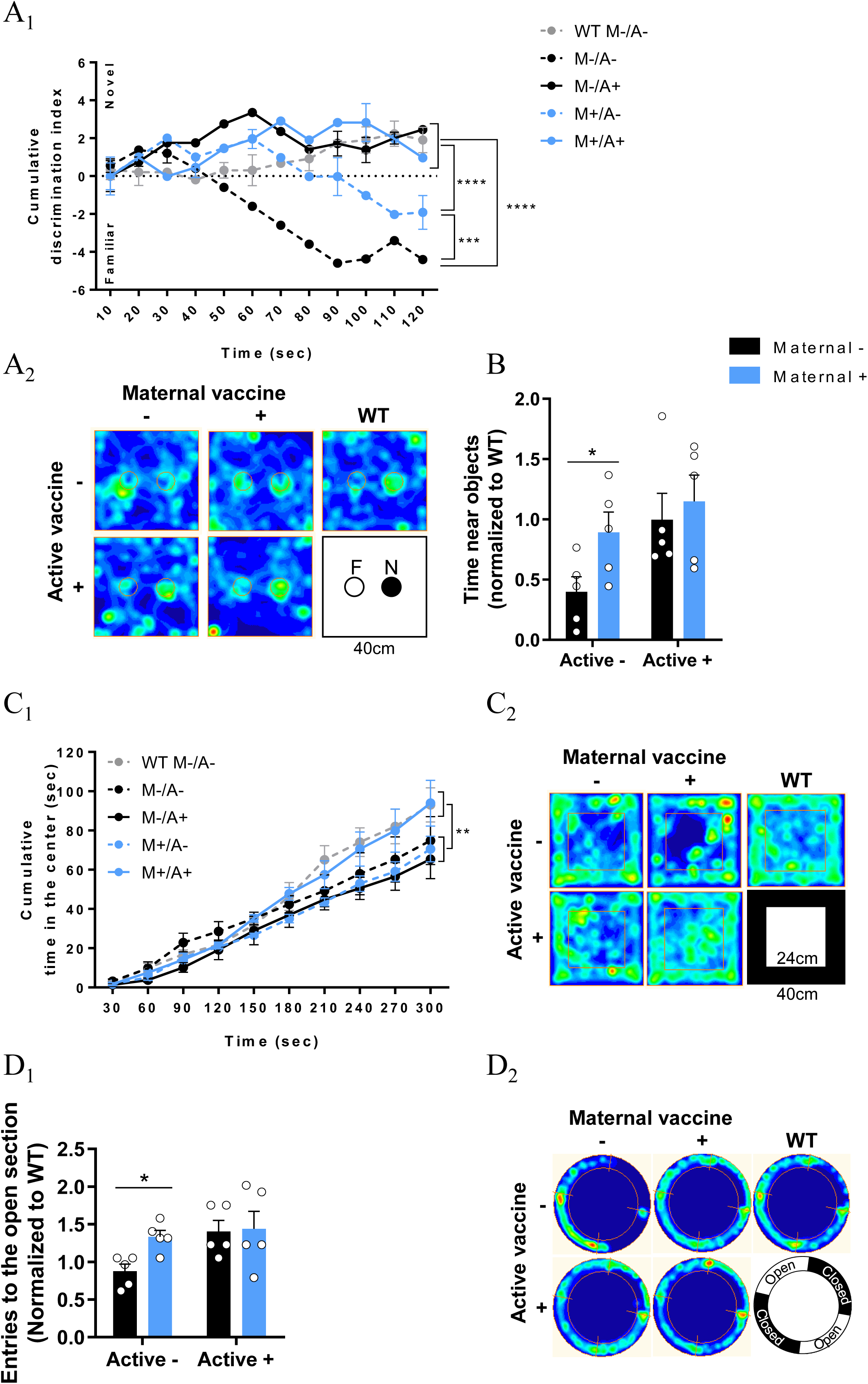
Combination of maternal and active vaccination rescues short-term memory abilities and normalizes exploratory behavior. A behavioral battery was conducted at 4m of age to assess the effect of maternal and active vaccinations on cognition. (A_1_) Short-term memory was assessed using the NOR test. (A_2_) Occupancy plots of NOR arena – F (familiar object), N (novel object). (B) Total time near objects in the NOR test. (C_1_) Exploratory behavior in the OF test. (C_2_) Occupancy plots of the OF arena. (D_1_) Anxiety-related behavior in the EZM. (D_2_) Occupancy plots of the EZM. *P<0.05, **P<0.01, ***P<0.001, ****P<0.0001, two-way ANOVA, repeated measures two-way ANOVA.

### Maternal vaccination reduces cerebral Aβ levels in adult 5xFAD mice

Mice were sacrificed at 5m to assess Aβ-related neuropathology (Fig. 1A). Cortical Aβ_40_ and Aβ_42_ levels were quantified using a modification of previously described sELISA (Illouz, Madar, Griffioen, & Okun, 2017) in soluble (TBST-soluble), pre-amyloid complexes (SDS-soluble) and insoluble-fibrillary (FA-soluble) Aβ extracts. M+/A- and M+/A+ mice exhibited reduced soluble Aβ_42_ compared with both M-/A- and M-/A+ mice (0±0, 0±0, 465.4±191.5, and 143±112.3pg/mg, respectively, P<0.05, Fig. 3A). No difference was observed between M-/A- and M-/A+ groups (P=0.16, Fig. 3A), although M-/A- was the only group to have elevated levels of soluble Aβ_42_ compared with WT controls (465.4±191.5, 0±0pg/mg, respectively, P<0.05, Fig. 3A). SDS-soluble pre-amyloid Aβ_42_ was reduced in M+/A- and M-/A+ but not in M+/A+ treated mice compared with M-/A-, (1.78±1.28, 2.47±1.43, 3.25±1.39, 8.33±2.13ng/mg, respectively, P<0.01, P=0.21, respectively, Fig. 3B). However, M-/A- mice were the only group to have elevated levels of SDS-soluble Aβ_42_ compared with WT controls (8.33±2.13, 0.54±0.34ng/mg, respectively, P<0.05, Fig. 3B). Importantly, insoluble Aβ_42_ was reduced in maternally vaccinated mice (M+/A- and M+/A+), compared with unvaccinated and actively vaccinated mice (M-/A- and M-/A+) (1063±249, 937±303.1, 2583±621, 2093±466ng/mg, respectively, P<0.01, Fig. 3C). No difference was observed in cortical insoluble Aβ_42_ between M-/A- and M-/A+ (P=0.68, Fig. 3C) or between M+/A- and M+/A+ (P=0.97, Fig. 3C). Elevated levels of insoluble Aβ_42_ were observed in M-/A- and M-/A+ compared with WT controls (2583±621, 2093±466, 31.24±24.82ng/mg, respectively, P<0.01, Fig. 3C). These findings suggest that active vaccination alone is insufficient in reducing insoluble Aβ_42_. Moreover, active vaccination did not yield additional reduction in already maternally immunized mice, suggesting that early passive maternal vaccination results in a long-lasting effect on either Aβ accumulation, clearance, or both.

**Fig. 3.**
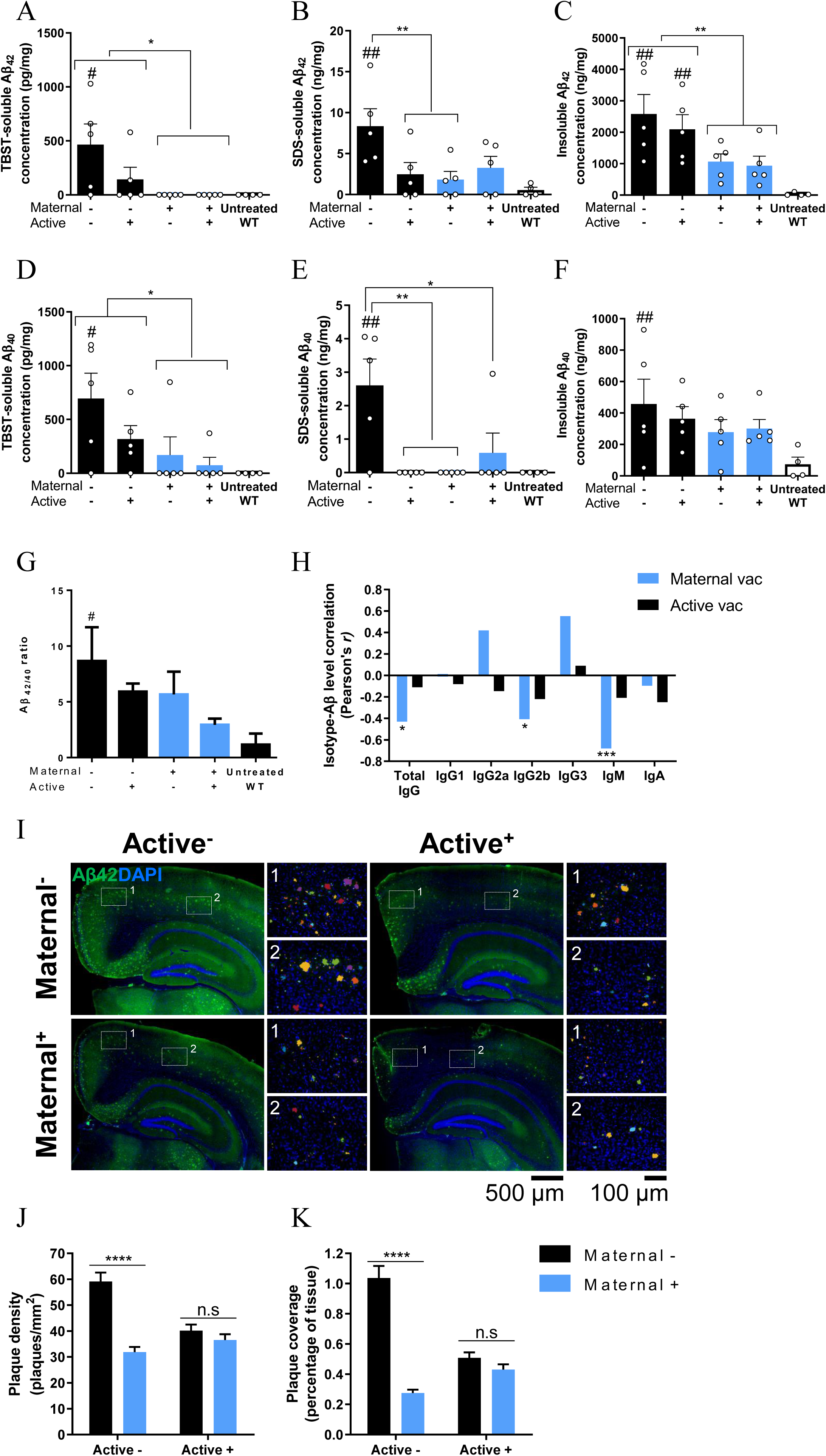
Maternal vaccination reduces Aβ pathology independently of anti-Aβ antibody presence at adulthood. Cortical Aβ levels were measured using sELISA (A) TBST, (B) SDS, and (C) formic-acid soluble Aβ_42_. (D) TBST, (E) SDS, and (F) formic-acid soluble levels of cortical Aβ_40_. (G) Cortical Aβ42/40 ratio. (H) Insoluble Aβ levels at 5m of age negatively correlate with IgG2b and IgM levels, following maternal vaccination. (I) Plaque load was quantified using immunofluorescence and blob detection. (J) Quantification of plaque density (plaques/mm^2^) and (K) Plaque tissue coverage (percentage), *P<0.05, **P<0.01, ****P<0.0001, #P<0.05, ##P<0.01 compared to WT controls, two-way ANOVA, one-way ANOVA, Pearson’s correlation.

Maternally vaccinated mice (M+/A- and M+/A+) exhibited reduced soluble Aβ_40_ compared with M-/A- and M-/A+ mice (169.4±169.4, 74.26±74.26, 695±236.1, 317.6±125.8pg/mg, respectively, P<0.05, Fig. 3D). M-/A+ mice exhibited a non-significant reduction in soluble Aβ_40_ compared with M-/A- mice (P=0.22, Fig. 3D). However, M-/A- was the only group in which soluble Aβ_40_ was significantly higher than WT controls (695±236.1, 0±0pg/mg, respectively, P<0.05, Fig. 3D). The SDS-soluble fraction of Aβ_40_ was lower in all vaccination groups compared with unvaccinated mice (0±0, 0±0, 0.59±0.59, 2.6±0.78ng/mg, respectively, P<0.01, P<0.01, P<0.05, respectively, Fig. 3E). Elevated levels of SDS-soluble Aβ_40_ were measured in M-/A- mice compared with WT controls (2.6±0.78, 0±0ng/mg, respectively, P<0.01, Fig. 3E). While M-/A- mice exhibited elevated levels of insoluble Aβ_40_ compared with WT controls (475.5±158.3, 73.45±46.37ng/mg, respectively, P<0.01, Fig. 3F), no difference was observed between 5xFAD groups (P=0.13, Fig. 3F). Insoluble Aβ_42_/Aβ_40_ ratio (Jarrett & Lansbury, 1993) was significantly higher in M-/A- mice compared with WT controls (8.67±3.02, 1.16±0.99, P<0.05, Fig. 3G), with no difference from all other 5xFAD groups (P=0.13, main effect maternal vaccination, P=0.16, main effect active vaccination, Fig. 3F). To test whether the reduction in cerebral Aβ among maternally vaccinated mice is a result of enhanced clearance or reduced production of Aβ, hAPP levels were measured at the transcript and protein levels. *hAPP* mRNA did not differ for both maternal (P=0.42) and active (P=0.35) vaccinations (Supplementary Fig. S3A), suggesting that reduced APP production cannot explain the observed reduction in Aβ. At the protein level, actively and combined vaccinated mice exhibited a reduction in hAPP compared with untreated and maternally vaccinated mice (0.76±0.65, 0.65±0.1 for M-/A+ and M+/A+, respectively, compared with 0.92±0.06, 1.01±0.07 for M-/A- and M+/A-, respectively, P<0.05, supplementary Fig. S3B). This effect accounts for only ∼18% of the final reduction seen in cerebral Aβ levels and is possibly due to neutralization of hAPP by antibodies. These results suggest that most of the reduction in Aβ levels is not due to lowered hAPP expression and is, therefore, attributable to Aβ clearance. Maternal anti-Aβ IgG levels in offspring circulation negatively correlated with insoluble Aβ_42_ levels at 5m of age (*r*=-0.43, P<0.05, Fig. 3H, S4A), whereas active vaccination showed no such correlation (*r*=-0.11, P=0.32, Fig. 3H, S4D). Similar correlations were found for maternal IgG2b and IgM levels (*r*=-0.4, *r*=-0.68, respectively, P<0.05, P<0.001, respectively, Fig. 3H, S4B, S4C), while active-vaccination-derived IgG2b and IgM did not correlate with reduction in Aβ load (*r*=-0.21, *r*=-0.1, respectively, P=0.17, P=0.18, respectively, Fig. 3H, S4E, S4F). M+/A- exhibited reduced plaque density in the cortex compared with M-/A- (31.9±1.98, 59.1±3.43 plaques/mm^2^, respectively, P<0.0001, Fig. 3I-J). Accordingly, plaque coverage percentage was also reduced among M+/A- compared with M-/A- (0.27±0.02, 1.03±0.08%, respectively, P<0.0001, Fig. 3I, K). Active vaccination had a minor contribution to plaque density reduction in already maternally vaccinated mice (40.15±2.32, 35.55±2.23 plaques/mm^2^, respectively, P=0.49, Fig. 3I-J).

Taken together, these findings provide evidence for a long-lasting effect of maternal vaccination during developmental and postnatal periods on Aβ clearance or accumulation, with a limited contribution of active vaccination at early adulthood.

### Maternal vaccination induces long-term FcRs upregulation in the brain

A limited-time exposure of offspring to maternal Ab resulted in a long-lasting effect on Aβ pathology and short-term memory restoration, months after Abs were not detected in the circulation. As Aβ was not targeted by maternal Abs later than 1m of age, two mechanisms could be ruled out: (a) maternal Abs interfere with oligomerization and fibrillation, to prevent plaque formation (Geylis & Steinitz, 2006); and (b) maternal vaccination facilitates Aβ clearance by microglia via opsonization (Strohmeyer et al., 2005). This led us to hypothesize that maternal vaccination promoted a long-lasting microglial phagocytic phenotype, independently of the presence of anti-Aβ Abs. To verify this, we assessed the expression of various Fc receptors in the brain at 1m following maternal vaccination only, and at 5m of age, following maternal and active vaccination. The activating FcγRI, FcγRIII, and FcγRIV initiate Fc-mediated phagocytosis via several actin-regulating pathways (Bruhns & Jonsson, 2015; Jaumouille et al., 2014). Additionally, FcR-neonatal (FcRn) is crucial for the transport of maternal Abs from the placenta and gut to the circulation of fetuses and newborns (Latvala, Jacobsen, Otteneder, Herrmann, & Kronenberg, 2017). At 1m, cerebral *FcγRI* levels were elevated in maternally vaccinated WT and 5xFAD mice compared with sham-vaccinated controls (1.56±0.16, 1±0.1 fold-change in WT, 1.77±0.16, 0.86±0.05 fold-change in 5xFAD, P<0.05, P<0.001, respectively, Fig. 4A). Interestingly, *FcγRIII* levels increased in maternally vaccinated 5xFAD mice compared with controls (1.6±0.16, 0.86±0.1 fold-change, respectively, P<0.001, Fig. 5B), but not among vaccinated WT mice compared with WT controls (P=0.12, Fig. 4B). A similar strain-specific effect was also observed for *FcγRIV* expression in the brain (P=0.07 for WT, 2±0.25, 0.54±0.045 fold-change, respectively, P<0.0001 for 5xFAD, Fig. 4C). *FcRn* levels were elevated in both WT and 5xFAD mice compared with controls (1.45±0.09, 1±0.054 fold-change for WT, respectively, P<0.01, 1.54±0.12, 1.08±0.07 fold-change, respectively, P<0.01 for 5xFAD, Fig. 4D). Interestingly, levels of the inhibitory *FcγRIIb* were elevated in vaccinated 5xFAD mice compared with controls (1.77±0.33, 0.91±0.09, fold-change, P<0.01, Fig. 4E), but not in vaccinated WT mice (P=0.18, Fig. 4E). These results indicate that the presence of maternal Abs correlated with transcript upregulation of their receptors in the brain. At 5m, *FcγRI* levels in M+/A-, M-/A+, and M+/A+ mice were elevated compared with M-/A- mice (1.44±0.11, 1.39±0.25, 1.44±0.2, 0.79±0.04 fold-change, respectively, P<0.05, Fig. 4F).

**Fig. 4.**
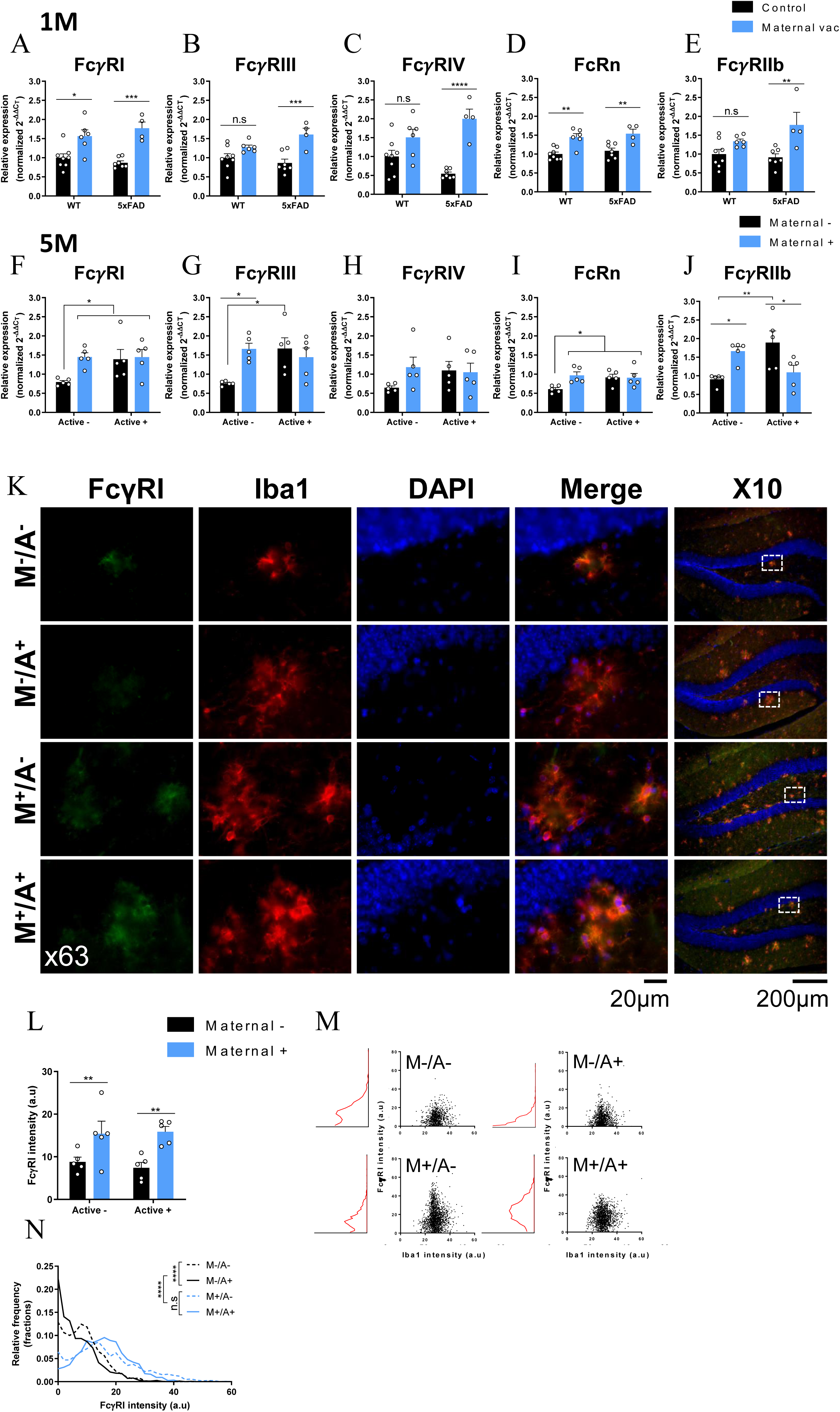
Maternal vaccination induces long-term FcRs upregulation in the brain. Levels of cerebral *FcR* mRNA and protein levels of FcγRI were quantified using RT-qPCR and immunofluorescence following maternal vaccination at 1m of age (A-E) and after maternal and active vaccination at 5m of age (F-N). mRNA levels of (A) *FcγRI*, (B) *FcγRIII*, (C) *FcγRIV*, (D) *FcRn*, and (E) *FcγRIIb* in 1m-old WT and 5xFAD mice. Levels of (F) *FcγRI*, (G) *FcγRIII*, (H) *FcγRIV*, (I) *FcRn*, and (J) *FcγRIIb* in maternally and actively vaccinated 5xFAD mice. (K) FcγRI expression was assessed using double-labeled immunofluorescence with Iba1^+^ microglia, using the x10 and x63 objectives for visualizing and quantification, respectively. (L) Quantification of FcγRI signal intensity among microglia. (M) Scatter plot and intensity distribution FcγRI signals in Iba1+ cells. (N) Overlay and comparisons of FcγRI expression distribution. *P<0.05, **P<0.01, ***P<0.001, ****P<0.0001, two-way ANOVA, Pearson’s correlation, corrected two-sample Kolmogorov-Smirnov test.

**Fig 5.**
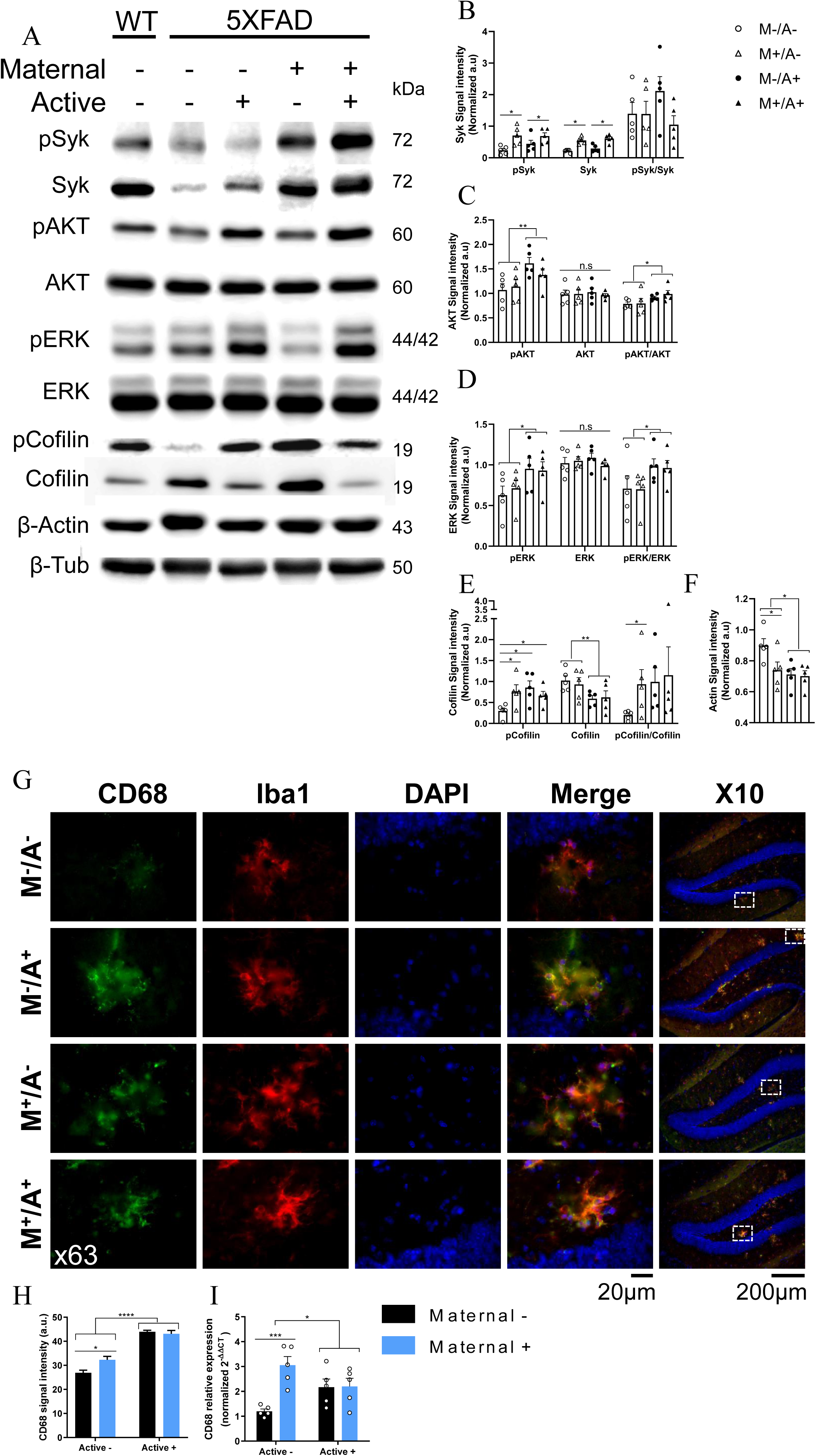
Maternal vaccination activates FcR-mediated phagocytosis via activation of AKT, ERK, and Cofilin actin-cytoskeleton regulation pathways. Level of Syk and downstream signaling molecules from the Fc-mediated phagocytosis pathway were measured using immunoblotting following maternal vaccination and active vaccination at 5m of age. Functionally, Fc-mediate Phagocytosis and hippocampal microglial reactivity were assessed using double labeling of CD68 and Iba1^+^ microglia. Levels of (A, B) pSyk/Syk, (A, C) pAKT/AKT, (A, D) pERK/ERK, (A, E) pCofilin/Cofilin, and (A, F) Actin levels, normalized to tubulin. (G) Double labeling of CD68 and Iba1^+^ microglia using the x10 and x63 objectives for visualizing and quantification, respectively. (H) Quantification of CD68 signal intensity. (I) CD68 mRNA levels were measured using RT-qPCR. *P<0.05, **P<0.01, ***P<0.001, two-way ANOVA.

No difference was observed between M+/A- and M+/A+ mice, implying the absence of additional contribution of active vaccination to *FcγRI* upregulation in already maternally vaccinated mice. *FcγRIII* mRNA levels were higher in M+/A- and M-/A+ mice compared with unvaccinated mice (1.62±0.14, 1.67±0.27, 0.75±0.02, respectively, P<0.05, Fig. 4G). However, combined maternal and active vaccination M+/A+ did not differ significantly from unvaccinated mice (1.44±0.24, 0.75±0.02, P=0.07, Fig. 4G). *FcγRIV* transcript levels did not differ between groups (P=0.27, main effect for maternal vaccination, P=0.47, main effect for active vaccination, Fig. 4H). *FcRn* levels in M+/A-, M+/A+, and M+/A+ mice were elevated compared with M-/A- mice (0.96±0.09, 0.91±0.08, 0.9±0.11, 0.6±0.04 fold- change, respectively, P<0.05, Fig. 4I). Levels of inhibitory *FcγRIIb* were elevated in M+/A- and M-/A+ compared with M-/A- (1.66±0.11, 1.89±0.31, 0.9±0.06, fold-change, P<0.05, P<0.01, respectively, Fig. 4J). Importantly, *FcγRIIb* levels were reduced in M+/A+ compared with M-/A+ mice (1.09±0.19, 1.89±0.31 fold-change, P<0.05, Fig. 4J), suggesting that the combined vaccination results in upregulation of activator FcRs and downregulation of inhibitory FcRs. Finally, elevation in *FcγRI, FcγRIII* and *FcRn* strongly correlated with Aβ reduction (full data are described in the supplementary text, Fig. S5).

### Maternal vaccination elevates FcγRI on microglial cells

As FcγR signaling could mediate the effects of maternal vaccination on Aβ pathology, we next assessed FcγRI, FcγRIIb, FcγRIII, and FcγRIV expression in CNS cells. FcγRI was abundantly expressed in microglia, but not in neurons or astrocytes (full data are described in the supplementary text, Fig. S6-7). Consequently, we next assessed FcγRI levels in microglial cells in the hippocampus and cortex of maternally and actively immunized mice. At 5m, hippocampal FcγRI levels were elevated in M+/A- and M+/A+ microglia compared with both M-/A- and M-/A+ (15.41± 6.58, 15.86±2.8, 8.9±2.8, 7.4±2.8 a.u (arbitrary unit), P<0.01, Fig. 4K-L). Active vaccination alone did not yield such elevation compared with M-/A-(P=0.84, Fig. 4K-L). FcγRI expression was normally distributed in M+/A+ microglia (P=0.7 compared with simulated normal distribution of the same mean and standard deviation, Fig. 4M) and close to normal distribution in M+/A- (P=0.04, Fig. 4M). In M-/A- and M-/A+, FcγRI distribution was skewed to the right and significantly differed from simulated normal distribution (P<0.0001, Fig. 4M). FcγRI distribution in M+/A- and M+/A+ hippocampal microglia differed from M-/A- and M-/A+ (P<0.0001, Fig. 4N), with no difference between M+/A- and M+/A+ mice, suggesting no additive effect of active vaccination on already maternally immunized animals. Similar effects were found in cortical microglia, as FcγRI levels were elevated in maternally vaccinated mice M+/A- and M+/A+ compared with controls (full data are described in the supplementary text, Fig. S8).

### Maternal vaccination activates phagocytosis-related signaling cascades in microglial cells

Maternal vaccination upregulated *FcγRI*, *FcγRIII*, *FcγRIV*, and *FcRn* transcripts (Fig. 4A-J) and increased the microglial FcγRI protein expression (Fig. 4K-N). Clustering of FcγRs induces activation of Src-related kinases that phosphorylate the immunoreceptor tyrosine-based activation motif (ITAM) domain on FcγRs. Syk is recruited to the receptor and undergoes autophosphorylation. This in turn activates downstream signaling molecules, such as AKT, ERK, and cofilin, all involved in actin-cytoskeleton regulation and are essential for phagocytosis (Kiefer et al., 1998).

At 5m of age, pSyk was upregulated in the hippocampus of M+/A- and M+/A+ compared with M-/A- and M-/A+ mice (0.68±0.13, 0.7±0.16, 0.23±0.07, 0.44±0.14, a.u, respectively, P<0.05, Fig. 5A-B). Similarly, total Syk was also upregulated in maternally vaccinated mice (0.6±0.05, 0.54±0.05, 0.21±0.02, 0.29±0.05, a.u, respectively, P<0.001, Fig. 5A-B), suggesting that FcγR upregulation by maternal vaccination resulted in activation of downstream Syk signaling. pAKT levels were elevated as a result of active vaccination in M-/A+ and M+/A+ compared with M-/A- and M+/A-, with no effect of maternal vaccination (1.64±0.12, 1.37±0.12, 1±0.13, 1.13±0.13, a.u respectively, P<0.01, Fig. 5A, C). Total AKT levels did not differ between groups (P=0.9, main effect for active vaccination, 0.76, main effect for maternal vaccination, Fig. 5A, C). In a similar manner, pERK levels increased in actively vaccinated mice M-/A+ and M+/A+ compared with M-/A- and M+/A-, with little contribution of maternal vaccination (0.95±0.13, 0.93±0.1, 0.62±0.11, 0.72±0.1 au, respectively, P<0.05, Fig. 5A, D). Total ERK levels did not differ between groups (P=0.98, main effect for active vaccination, 0.54, main effect for maternal vaccination, Fig. 5A, D). This may serve as evidence for Syk activation of ERK and AKT- mediated actin regulation as a result of active, but not maternal, vaccination. pCofilin levels increased in maternally immunized mice M+/A-, compared with unvaccinated mice M-/A- (0.76±0.2, 0.29±0.07 a.u, respectively, P<0.05, Fig. 5A, E). No difference was observed between actively M-/A+ and combined M+/A+ vaccination (P=0.87, Fig. 5A, E). Moreover, maternal vaccination alone yielded similar pCofilin levels as did active vaccination or combined vaccine. Levels of cofilin were reduced in actively M-/A+ and combined M+/A+ mice compared with unvaccinated M-/A- and maternally vaccinated M-/A+ mice (0.59±0.07, 0.62±0.15, 1.02±0.1, 0.93±0.16 au, respectively, P<0.05, Fig. 5A, E). This may serve as evidence that maternal vaccination enhances actin polymerization via Syk-mediated elevation in pCofilin levels and reduction in Cofilin levels. Finally, we found that both maternal M+/A-, M+/A+, and active M-/A+ vaccination reduce actin levels compared with unvaccinated M-/A- mice (0.73±0.05, 0.7±0.03, 0.71±0.03, 0.89±0.04, a.u, respectively, P<0.05, Fig. 5A, F). These findings provide evidence for the long-term effect of maternal vaccination in facilitating FcR-mediated phagocytosis signaling pathways (Fig. S9A).

### Maternal vaccination facilitates long-term Aβ clearance via enhancing FcR-mediated microglial phagocytosis

FcR upregulation led to recruitment and activation of Syk, which in turn initiated an actin regulating pathway via AKT, ERK, and Cofilin. Regulation of actin polymerization and depolymerization is essential for the construction of the phagosome (Jaumouille et al., 2014). CD68, a lysosomal protein expressed in macrophages and activated microglia, is often associated with pro-inflammatory disease-associated microglia. To assess whether maternally and actively vaccinated mice exhibit elevated phagocytic behavior, immunohistochemical (IHC) analysis of CD68 among Iba1^+^ cells was conducted. Maternally vaccinated mice M+/A- exhibit elevated CD68 levels in Iba1^+^ cells, compared with M-/A- (32.2±1.4, 26.98±1.07 a.u, respectively, P<0.05, Fig. 5G-H). Moreover, M-/A+ and M+/A+ mice exhibit higher CD68 expression than M-/A- and M+/A- mice (43.9±0.6, 43.09±1.4 a.u, respectively, P<0.0001, Fig. 5G-H), with no difference between M-/A+ and M+/A+ (P=0.75, Fig. 5G-H). These findings suggest that maternal vaccination alone enhances microglial phagocytosis. At the transcript level, M+/A- mice exhibit elevated levels of *CD68* mRNA compared with M-/A- (3.05±0.34, 1.19±0.09 fold-change, respectively, P<0.001, Fig. 5I), suggesting that maternal vaccination support microglial phagocytosis even in the absence of Abs, possibly via FcR-Syk activation. Additionally, actively vaccinated M-/A+ and combined M+/A+ vaccination mice also exhibited elevated transcript levels of *CD68* compared with unvaccinated controls (2.16±0.3, 2.19±0.3 fold-change, respectively, P<0.05, Fig. 5I). This data imply that microglial phagocytosis is enhanced in maternally and actively vaccinated mice. To support the maternal vaccination/FcR/Syk/phagocytosis hypothesis, we stained hippocampal CD68^+^ cells for pSyk. Maternally vaccinated M+/A- mice exhibited elevated pSyk levels among CD68^+^ cells in the hippocampus, compared with M-/A- mice (18.68±0.51, 16.57±0.52 a.u, respectively, P<0.05, Fig. S10A-B), suggesting that maternal vaccination alone is sufficient in inducing Syk activation in microglial cells, long after maternally-induced Abs are abolished. Additionally, maternally and actively M+/A+ mice exhibited elevated microglial pSyk levels compared with actively M-/A+ immunized mice (18.87±0.37, 14.09±0.39 a.u, respectively, P<0.0001, Fig. S10A-B), suggesting that active vaccination alone does not not induce Syk activation as strongly as maternal vaccination.

### Maternal antibodies facilitate Aβ clearance by microglial cells in a Syk-dependent manner

We next assessed the effect of maternal antibodies on Syk-dependent phagocytic capacity of N9 murine embryonic microglial cell-line (Corradin, Mauel, Donini, Quattrocchi, & Ricciardi-Castagnoli, 1993) and adult primary microglia. Increasing concentration (0.75-5µM) of BAY-61-3606 (BAY), a highly selective Syk inhibitor (Yamamoto et al., 2003), in the media of N9 cells decreased Aβ-induced Syk phosphorylation in a dose-dependent manner (Fig. 6A). Accordingly, alteration in downstream signaling molecules of the Fc-mediate phagocytosis pathway was observed, namely, reduction in ERK and AKT phosphorylation and an increase in cofilin phosphorylation (Supplementary Fig. S11A-C). As activation of ERK and AKT exhibited a considerable reduction in cells incubated with 5µM of BAY, the following experiments were conducted using this concentration. We first assessed the general capacity of N9 cells to perform phagocytosis of FBS-coated fluorescent beads following a 3h FBS deprivation. N9 cells treated with BAY exhibited reduced phagocytic activity compared with controls (8.92±0.16, 23.97±1.12 percentage of FITC+ cells, respectively, P<0.001, Fig. 6B-C) and a reduction in FITC intensity (8958±156.42, 9535±326.64, mean FITC intensity, a.u, respectively, P<0.05, Fig. 6D), reflecting not only a lower percentage of phagocytic cells, but also a lower number of intracellular beads. Importantly, Syk inhibition reduced phagocytosis of Aβ_42_ peptide, as the number of Aβ foci per mm^2^ was reduced by ∼50% following BAY administration compared to control (48.32±4.18, 96.38±6.71 Aβ foci per mm^2^, respectively, P<0.0001, Fig. 6E-F). Next, we tested whether pre-incubation of Aβ with serum from anti-Aβ vaccinated dams facilitates phagocytosis of Ab-Ag complexes. N9 cells were treated with Aβ_42_ alone, Aβ_42_ pre-incubated with serum from sham, or anti- Aβ vaccinated dams with BAY. 5.1% of untreated N9 cells were Aβ positive, demonstrating baseline phagocytosis capacity of these cells (Fig. 6G-H). Incubation with serum from sham-vaccinated resulted in 11.7±0.03% and of phagocytic cells (P<0.0001 compared to baseline, Fig. 6G-H), and incubation with serum from anti-Aβ vaccinated dams elicited phagocytosis in 81±0.003% of the cells (P<0.0001 compared with baseline and sham-vaccinated serum, Fig. 6G-H). Importantly, this drastic effect was abrogated by Syk inhibition, as only 18.3±0.003% of BAY and serum-treated cells were phagocytic (P<0.0001, Fig. 6G-H). The median of FITC intensity increased among cells treated with anti-Aβ serum compared with sham-vaccinated serum (316±96.1, 168±30.4 a.u, respectively, P<0.0001, Fig. 6I-J) and reduced following treatment with BAY (183±30.7 a.u, P<0.0001, Fig. 6I-J). These results demonstrate that maternal antibodies facilitate Fc-receptor mediated Aβ phagocytosis in a Syk-dependent pathway. To further understand the alteration in cellular signaling following serum and Aβ administration, Syk phosphorylation was measured using western blotting 10-30min after treatment with either Aβ_42_ alone, serum from anti-Aβ vaccinated dams, or Aβ_42_ pre-incubated with serum from vaccinated and sham-vaccinated dams (Fig. 6K). Aβ alone elicited elevation in Syk activation (3.75-fold change of naïve cells after 20min, Fig. 6K). Serum from anti-Aβ vaccinated dams alone also resulted in Syk activation (4.06-fold change of naïve cells after 10min, Fig. 6K). The difference in Syk activation timing implies that Aβ and antibodies alone may activate different signaling pathways, e.g. Fc-receptor signaling by antibodies and Trem2 signaling by Aβ, both of which are Syk-dependent. Interestingly, serum from sham-vaccinated dams elicited a milder activation of Syk (1.92-fold change of naïve cells after 10min, Fig. 6K), supporting our hypothesis that Syk phosphorylation associated with Fc-receptors peaks 10min following treatment. Treatment with both Aβ and serum from vaccinated dams yielded a strong but transient Syk activation (4.16-fold change of naïve cells after 20min, Fig. 6K), suggesting that Ab-Ag complexes generate a somewhat different effect than Ab or Ag alone.

**Fig. 6.**
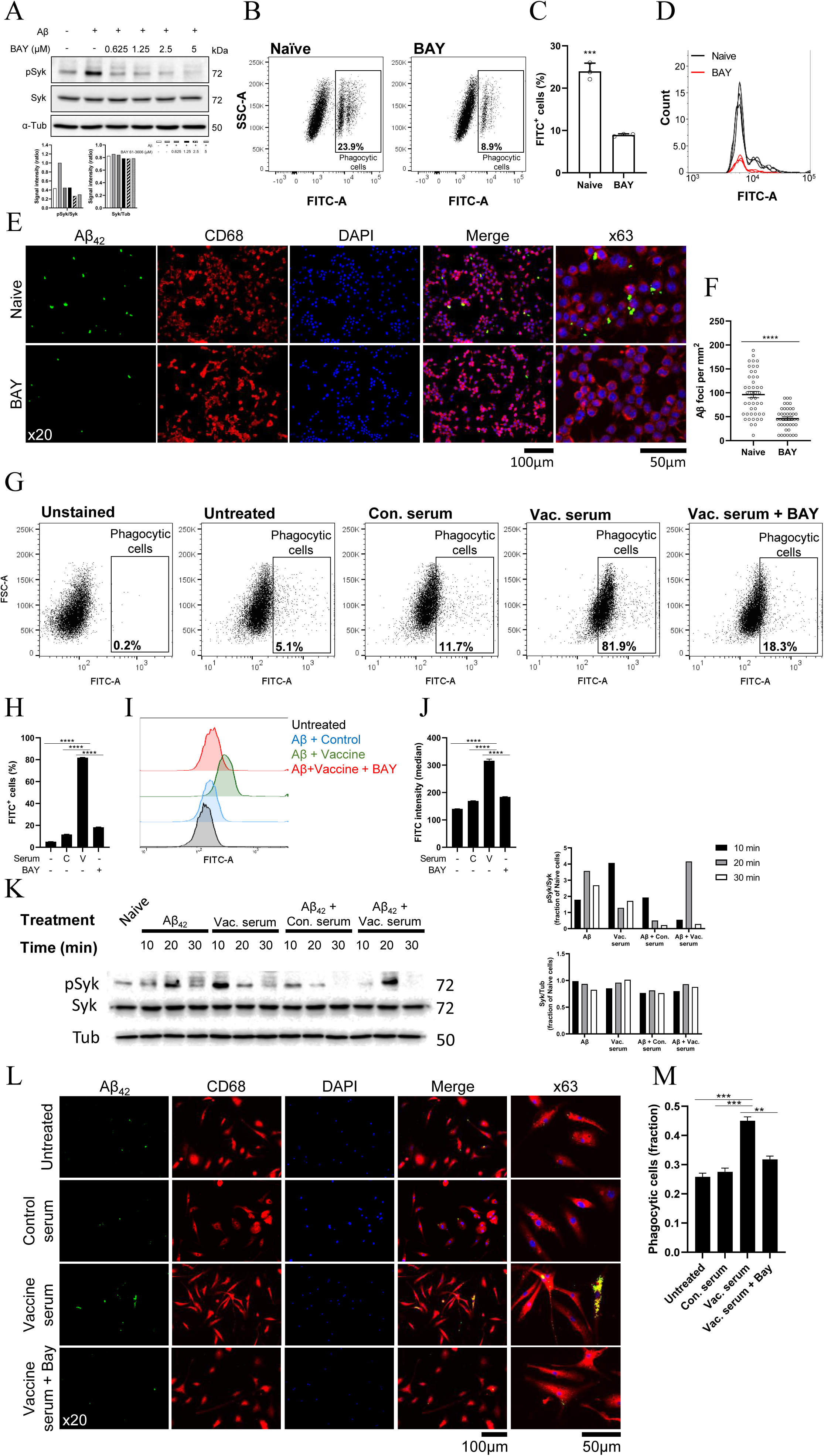
Maternal antibodies mediate Aβ phagocytosis in N9 microglial cell-line and primary adult microglia in a Syk-dependent manner. Phagocytosis capacity of N9 murine embryonic cell-line and adult primary microglia, as well as intracellular signaling, were assessed following treatment with either Aβ_42_ alone or Aβ_42_ pre-incubated with serum from anti-Aβ or sham-vaccinated dams. The contribution of Syk signaling was assessed using Syk inhibition by BAY-61-3606. (A) BAY inhibits Syk phosphorylation in a dose-dependent manner. (B-D) Phagocytosis of FBS-coated fluorescent beads by N9 cells is reduced following treatment with Syk inhibitor. (E-F) IF of N9 cells following incubation with Aβ reveals a reduction in phagocytosis capacity after Syk inhibition. (G-J) Serum from anti-Aβ vaccinated dams increases phagocytosis by N9 cells compared with serum from sham-vaccinated dams and abrogation of this effect following Syk inhibition. (K) Treatment with Aβ, serum from anti-Aβ or sham-vaccinated dams, or combined treatment reveals differences in the time-course of Syk phosphorylation. (L-M) Phagocytosis assay of Aβ by adult primary microglia reveals that serum from anti-Aβ vaccinated dams facilitates Aβ clearance in a Syk-dependent pathway. **P<0.01, ***P<0.001, ****P<0.0001, unpaired t-test, Chi-squared test for independence, Fisher’s exact test, one-way ANOVA.

To further examine the contribution of maternal antibodies to Fc-receptor Syk-mediated Aβ clearance by microglia, phagocytosis assay was conducted using adult primary microglia from naïve WT mice. Cells were administered with Aβ_42_ (750 nM) following incubation with serum from sham-vaccinated dams, anti-Aβ vaccinated dams, and Syk inhibitor BAY. 4h following Aβ administration, primary microglial cells were fixed and stained for CD68 and intracellular Aβ. Treatment with serum from sham-vaccinated females did not increase Aβ phagocytosis compared with untreated cells (0.27±0.07, 0.25±0.07, a fraction of Aβ+ cells, respectively, P=0.37, Fig. 6L). In contrast, treatment with anti-Aβ serum increased phagocytic activity compared with both untreated and sham-vaccinated serum treatment (0.45±0.08 fraction of Aβ+ cells, P<0.001, Fig. 6L). Importantly, this effect was abrogated by Syk inhibition (0.31±0.07 fraction of Aβ+ cells, P<0.01, Fig. 6L), demonstrating that Aβ clearance facilitated by antibodies is a Syk-dependent process.

## Discussion

To date, there is no effective therapy for AD-related dementia in individuals with DS. Considering the positive outcomes of active vaccination in Ts65Dn mouse model of DS (Illouz et al., 2019) and the rapid propagation of cerebral Aβ in DS, maternal vaccination may serve as early, preventative intervention, well before the formation of Aβ plaques. So far, anti-Aβ vaccination targeting dementia in AD patients has failed to translate to the clinic. However, the age of intervention in relation to the onset of the disease and to the subjects’ immunological age might be held responsible (Miller, 2012; Morris et al., 2012; Reiman et al., 2011). The DS case is therefore quite different in this manner. DS can be detectable in utero, and the progression of Aβ pathology is earlier and faster than in late-onset AD. We thus propose that early anti-Aβ intervention in the form of maternally transferred Abs, along with active postnatal vaccination, may provide continuous immune targeting of Aβ in individuals with DS.

In this study, we vaccinated WT females prior to their pregnancy with transgenic embryos, as the duration of mature IgG production exceeds the length of pregnancy in mice. In humans, a full humoral response can be achieved during pregnancy. In contrast to the full Aβ_42_, using the 1-11 fragment of Aβ, which does not contain cytotoxic T-cell epitopes, reduces the risk of a hazardous response (Olkhanud et al., 2012). Moreover, WT mice that received anti-Aβ_1-11_ vaccine did not exhibit any behavioral or cognitive decline, or neuronal loss, or inflammatory response compared with controls (Illouz et al., 2019). To date, more than 50 clinical trials have been conducted using DNA delivery by electroporation, reporting enhancement of DNA immunogenicity in humans and tolerability by patients without causing negative side effects (Rosenberg, Fu, & Lambracht-Washington, 2018).

The transfer of IgG2b Abs from dams to pups is supported by elevation of FcRn expression, which facilitates the passage of maternal Abs through the placenta and the crossing of maternal Abs from the gut of newborns into the circulation (Latvala et al., 2017). Moreover, FcRn plays a role in elongating IgG half-live as it restores opsonized IgG and prevents its degradation. Maternal vaccination thus initiates a positive regulation loop in which the presence of maternal Abs elevates the transcription of FcRn, which in turn supports the passage of these Abs through the placenta and lactation and elongates their half-live and functionality. Among the maternal Ab milieu, IgG2b was the most common isotype that crossed the placenta. IgG2b and IgM were the most common isotypes delivered through lactation. Since IgM cannot cross the blood-brain barrier (BBB) (Nilaratanakul et al., 2018), IgG2b is the main candidate effector for any downstream immunomodulation inside the CNS. Accordingly, we found elevated levels of *FcγRI* and *FcγRIII*, which bind IgG2b (Bruhns & Jonsson, 2015), but no expression of cerebral *Fcα/μR* that binds IgA and IgM, respectively. The positive correlation between maternal and newborns IgG2b levels along with the negative correlation between IgG2b levels in newborns and their Aβ levels at 5m strongly support this suggestion. Indeed, we found that high maternal IgM levels in newborns were correlated with low Aβ pathology at 5m. The contribution of IgM, if exists, is thus attributed to peripheral rather than CNS immunomodulation. Following active vaccination, IgG2b was the only isotype to remain high three months after the last boost. Taken together, IgG2b remains the main effector of both maternal and active vaccination. We hypothesized that a combination of maternal and active vaccination would yield continuous protection from Aβ neurotoxicity. Indeed, this combination resulted in normalization of exploratory behavior and restoration of short-term memory capacity. Maternal vaccination alone had no effect on exploratory behavior and only a partially positive effect on short-term memory. In contrast, active vaccination yielded a full rescue of short-term memory ability. Reduction in Aβ levels, however, was strongly dependent on maternal vaccination. Active vaccination produced a minor reduction in insoluble Aβ levels, whereas maternal vaccination, with or without the addition of active vaccination, greatly reduced levels of this fraction. Moreover, the newborn serum concentration of maternal Abs was significantly more predictive of Aβ clearance than actively-induced-Abs in young adults. Taken together, the cognitive and neuropathological findings indicate that the combination of maternal and active vaccination produced the most potent therapeutic effect.

Maternally induced Abs were present in the circulating blood of offspring for the short and limited period of pregnancy and lactation. Following weaning, these Abs were no longer detectable. However, four months after Abs were absent, Aβ levels were dramatically reduced, indicating a long-lasting immunomodulatory effect on microglial cells. Maternal vaccination indeed increased the expression of effector *FcγR*s and *FcRn* in the brain. Importantly, the expression levels of Fc receptors negatively correlated with insoluble Aβ levels. Fc receptors, expressed on various immune cells, constitute critical elements for activating or downregulating immune responses (Takai, 2005). Our data imply that an increase in FcR ligands upregulates the expression of *FcγRI*, *FcγRIII*, and *FcγRIV* at the transcript level in the brain, and FcγRI at the protein level in microglial cells. The effector FcγRI contains an intracellular immunoreceptor tyrosine-based activation motif ITAM domain, which mediates cellular activation triggered by Ab-receptor binding (Takai, 2005). Clustering of FcγRs induces activation of Src family kinases that phosphorylates the ITAM domain within the receptor. When an effector FcR is activated, Syk is recruited to the receptor and undergoes autophosphorylation, resulting in activation of downstream signaling molecules, such as AKT, ERK, and cofilin, all involved in actin-cytoskeleton regulation and are essential for phagocytosis (Kiefer et al., 1998). The FcγRs-Syk pathway is heavily implicated in phagocytosis (Song, Tanaka, Cox, & Lee, 2004; Stangel & Compston, 2001), a central function of microglial cells in the context of targeting amyloid burden (Krasemann et al., 2017). Indeed, we found that Syk was activated in maternally vaccinated mice. Accordingly, levels of phospho-cofilin were increased, and levels of cofilin decreased. The actin-depolymerizing factor (ADF)/cofilin protein family consists of small actin-binding proteins that play central roles in accelerating actin turnover by disassembling actin filaments (Tanaka et al., 2018). The activity of cofilin is regulated by phosphorylation on residue Ser3 by LIM kinases (LIMK1 and LIMK2) and TES kinases (TESK1 and TESK2), which inhibit its interaction with actin (Kanellos & Frame, 2016). Thus, maternal vaccination-induced FcR-Syk mediated cofilin phosphorylation contributes to the inactivation of cofilin and promotes actin polymerization, which is essential for phagosome formation in microglia (Jaumouille et al., 2014). ERK and AKT activation is also implicated in Fc-mediated phagocytosis, as both molecules undergo phosphorylation under Syk activation. Inhibition of Src and Syk kinases suppresses phagocytosis in primary human microglia with suppression of ERK and AKT activity (Song et al., 2004). Following the onset of Aβ propagation, activation of ERK and AKT increased in actively, but not maternally, vaccinated mice.

CD68 is a heavily glycosylated type I trans-membrane glycoprotein, mainly associated with endosomal/lysosomal compartments (Chistiakov, Killingsworth, Myasoedova, Orekhov, & Bobryshev, 2017). In the brain, it is highly expressed among phagocytic microglia (Fu, Shen, Xu, Luo, & Tang, 2014). In accordance with previous results, maternally vaccinated mice express higher levels of microglial CD68, suggesting enhanced phagocytic activity.

In line with these data, we found that N9 and primary microglia phagocytosis of Aβ is dramatically increased in the presence of anti-Aβ antibodies obtained from vaccinated dams. This effect is antibody-specific, as serum from sham-vaccinated dams (containing high titers of anti-HBSAg Abs) did not yield an equivalent increase. Importantly, antibody-mediated phagocytosis is a Syk-dependent process, as inhibition of Syk abrogated phagocytic increases by antibodies in both N9 and primary cells. Taken together, the association of Aβ clearance with microglial Syk activation maternal vaccination *in vivo* and the establishment of causal relations between maternal antibodies and phagocytosis capacity *in vitro* serve as evidence for the modulatory effect of maternal antibodies on microglial cellular function in eliminating Aβ from the brain parenchyma.

Taken together, we propose that during a short and limited period, maternal vaccination had set the infrastructure for facilitating future phagocytosis and clearance of Aβ by microglia. Such infrastructure consists of upregulation of effector FcγR in the brain. This Abs-induced process is supported by a positive regulation loop of Abs-FcRn that facilitates Ab delivery via the placenta and lactation and increases Abs functionality. Upon Aβ- expression, this infrastructure facilitates Fc-mediated phagocytosis and Aβ clearance, without the addition of active vaccination.

## Conclusion

In this study, we tested the efficacy of maternal vaccination in reducing Aβ-related neuropathology and cognitive decline. We found that maternal vaccination reduced dramatically Aβ pathology independently of active vaccination, in a long-lasting manner, months after maternal Abs were undetectable. A combination of maternal and active vaccination yielded the most potent results in ameliorating cognitive decline. Mechanistically, we propose that maternal vaccination activates the FcR-Syk-Cofilin axis, resulting in actin regulation and promotion of Aβ clearance by microglial cells. *In-vitro* phagocytosis was indeed facilitated by maternal antibodies, via Syk-dependent pathways. Maternal vaccination may thus provide a novel therapeutic approach for preventing early Aβ accumulation and dementia in DS individuals.

## Materials and Methods

### Study design

8-week old C57BL6 WT female mice (n=12/group) were actively vaccinated using the AβCoreS DNA-vaccine (Fig. 1A). Females were then crossed with 5xFAD males. IgG transfer via the placenta and lactation was assessed in one cohort (n=8/group), while long-term effects of maternal and active vaccination were tested in a second cohort (n=5/group). At 1m, active vaccination against Aβ or sham was administered to maternally or sham vaccinated offspring, to yield four experimental groups: sham-vaccinated M-/A-, maternally vaccinated M+/A-, actively vaccinated M-/A+, and combined maternal and actively vaccinated mice M+/A+. Sham-vaccinated WT mice were used as healthy controls.

### Animals

The 5xFAD EOAD mouse model (Jackson Laboratories #34840), which encompasses five AD-related mutations within the *APP* and *PSE1* genes (Oakley et al., 2006), was used in this study to model early Aβ accumulation in DS. 5xFAD mice were generously provided by Michal Schwartz (The Weizmann Institute, Rehovot, Israel). C57BL mice were used as dams and healthy controls (Jackson Laboratories #000664, Bar Harbor, ME). Animal care and experimental procedures followed the NIH Guide for the Care and Use of Laboratory Animals and Bar-Ilan and were approved by the Bar-Ilan University Animal Care and Use Committee.

### AβCoreS vaccine

AβCoreS is based on the pVAX1 expression vector (Olkhanud et al., ^2^_01_^2^) and encodes the N-terminus-Aβ_1-11_ fused to a Hepatitis-B surface antigen (HBsAg) and the Hepatitis-B core antigen (HBcAg), which acts to facilitate Ab production (Pyrski et al., 2017). As a control treatment, an expression vector (pUC19, New England Biolabs) containing HBsAg was used.

### Vaccine administration

Mice were intramuscularly injected at three time-points (14-day intervals), with 25µg DNA (50µl), as previously described (Olkhanud et al., 2012). Full electroporation configurations are detailed in the Supplementary Materials section.

### Antibody titer

Anti-Aβ1-11 Ab serum levels were quantified using a standard indirect ELISA against recombinant Aβ1-11 peptide. A detailed protocol is found in the Supplementary Materials section.

### Exploratory behavior

Exploratory behavior was recorded using a 40×40cm open field (OF) arena. The outer 8cm were defined as the area periphery and the 24×24cm inner square as the center. Illumination was kept at 1300lux. Mice were allowed to freely explore the arena for 5min (Seibenhener & Wooten, 2015). Analysis of animal behavior in this task, as well as the following tasks, was conducted using Anymaze (Stoelting).

### Anxiety assessment

Anxiety-related behavior was monitored using the elevated zero maze (EZM), a ring-shaped 65cm-high table, divided into closed and opened sections. The ring is 7cm wide and has an outer diameter of 60cm. The closed sections are confined by 20cm-high walls and a semi-transparent ceiling, whereas the opened sections have a 0.5cm high curbs at the edges. Illumination was kept at 1300lux, and trial duration was 5 minutes (Shepherd et al., 1994).

### Short-term object recognition memory

Short-term memory was assessed using the novel object recognition (NOR) test as previously described (Leger et al., 2013; Lueptow, 2017). Briefly, mice were placed in a 40×40cm arena with two different objects. In an acquisition trial, mice were allowed to explore their environment. In the following test trial, one of the objects was replaced by a novel object. Time spent near each of the objects was measured, and a preference index was calculated as the difference between time spent in the novel and familiar objects, divided by the total time spent near both objects.

### Spatial short-term memory

We utilized a variant of the T-maze alternation test, as previously described (Deacon & Rawlins, 2006). Full apparatus settings are found in the supplementary materials section.

### Brain Sample collection

Mice were anesthetized using Ketamine-Xylazine (100mg/kg, Vetoquinol, France, 10mg/kg, Eurovet, The Netherlands, respectively) and perfused with PBS. For Histology, hemibrains were transferred to 4% paraformaldehyde (PFA) at 4^°^C for 48h. Following fixation, tissues were transferred to a gradient of 20% and 30% sucrose aqueous solutions for 24h each. Hemibrains were then dissected into 40μm-thick slices using a microtome and stored in a cryoprotectant solution (30% glycerol and 35% ethylene glycol) at -20^°^C. For biochemical analysis, the cerebral cortex and hippocampi were separated, frozen on dry ice, and stored at −80^°^C.

### Measuring Aβ_40/42_ levels using sELISA

Aβ_40_ and Aβ_42_ in the cortex were measured using a modification of a previously published sandwich-ELISA protocol (Illouz et al., 2017). The full protocol is found in the Supplementary Materials section.

### Immunofluorescence

40μm-thick hemibrains were rinsed 5 times in 0.1% PBS-Triton for 5min. Nonspecific bindings were blocked using 20% normal horse serum in PBS-T for 1h at RT. For Aβ staining, antigen retrieval was conducted using incubation with 75% formic acid for 2min at RT. For pSyk, antigen retrieval was conducted using sodium citrate buffer (pH=6) for 20min at 95°C. Primary Abs for the following antigen were applied and incubated overnight at 4°C: Aβ_42_, Iba1, NeuN, GFAP, FcγRI, FcγRIIb, FcγRIII, FcγRIV, CD68, and pSyk. Next, sections were rinsed five times in PBS-T for 5min, and fluorescence-tagged secondary Abs were applied for 1h at RT. Primary and secondary Ab details are found in Supplementary Table 1. Slices were then stained with Hoechst 33342 (H3570, Invitrogen, Carlsbad, CA) and diluted at 1:1,000, followed by five 5-minute rinses with PBS-T.

### Aβ Plaque quantification

Cerebral Aβ plaques were quantified computationally using blob detection tools in MATLAB (MathWorks, Natick, MA). Briefly, montage immunofluorescence images of the hippocampus and cortex were obtained using the X10 objective of a Leica DM6000 microscope (Leica Microsystems, Wetzlar, Germany), coupled to a controller module and a high sensitivity 3CCD video camera system (MBF Biosciences, VT) and an Intel Xeon workstation (Intel). Automated imaging was implemented using the Stereo Investigator software package (MBF Biosciences, VT). Analyzed brain sections spanned from -1.355mm to -2.88mm from Bregma. A total of six sections, every 7th to 8th section (280–320μm apart), were stained, imaged, and fed to a pre-validated plaque detection algorithm. Automated plaque quantification was optimized and validated prior to use by comparing the results of a validation set to the results from three independent human countings.

### Quantification of microglial markers

Hippocampus and cortex were outlined according to the Paxinos atlas of the mouse brain. An average of 15 sections, spanning from 1.055mm to -3.68mm from Bregma, were used. Single-cell Iba1, FcγRI, CD68, and pSyk fluorescence intensity were filtered for noise reduction and calculated using MATLAB (MathWorks, Natick, MA).

### RT-qPCR

Cerebral Gene-expression was measured using standard TRIzol RNA extraction, cDNA generation, and RT-qPCR protocols. A detailed method is found in the Supplementary Materials section. Primers used are detailed in Supplementary Table 1.

### Western blot

A standard western blotting protocol was applied to measure hippocampal protein and phospho-protein. A detailed protocol is found in the Supplementary Materials section. Primary and secondary Ab details are found in Supplementary Table 1.

### N9 microglial cell-line culture

Murine embryonic microglia cell-line N9 (Corradin et al., 1993) were grown in Dulbecco’s modified Eagle’s medium (DMEM), supplemented with 10% fetal bovine serum (FBS), penicillin, streptomycin, and L-glutamine for 3-4 days to reach confluence.

### Murine primary microglia culture

Adult primary microglia were harvested from 2- month old C57BL6/j mice and cultured, as previously described (Moussaud & Draheim, 2010). Briefly, mice were anesthetized using Ketamine-Xylazine (100mg/kg, Vetoquinol, France, 10mg/kg, Eurovet, The Netherlands, respectively) and perfused with cold Hanks’ Balanced Salt Solution (HBSS, 14175, Thermo-Fisher Scientific, Waltham, MA). Brains were minced with a scalpel in an enzymatic solution containing papain and were incubated for 90 minutes at 37°C, 5% CO_2_. After 90 minutes, the enzymatic reaction was quenched with 20% FBS in HBSS, and cells were centrifuged for 7 minutes at 200g. The pellet was resuspended in 2ml of 0.5 mg/ml DNase in HBSS and incubated for 5min at room temperature. The tissue was gently disrupted, filtered through a 70mm-cell strainer, and centrifuged at 200g for 7 minutes. The pellet was resuspended in 20ml of 20% isotonic Percoll in HBSS, and pure HBSS was carefully overlaid on top of the cells-Percoll layer, followed by centrifuging at 200g for 20 min with slow acceleration and no brakes. The pellet, containing the mixed glial cell population, was washed once in HBSS and suspended in Dulbecco’s Modified Eagle’s/F12 medium (DMEM/F12) supplemented with 10% FBS, penicillin, streptomycin, L-glutamine and 10 ng/ml of carrier-free murine recombinant granulocyte and macrophage colony-stimulating factor (GM-CSF, 415-ML, R&D systems, Minneapolis, MN). Cell suspension from one brain was plated in a T75-cell culture flask coated with poly-D-lysine (P7280, Sigma, St. Louis, MO) and maintained at 37°C, 5% CO_2_ humidified incubator. The medium was changed twice a week. When full confluency was reached (after ∼2 weeks), the floating cell layer, which includes pure microglia, was collected and centrifuged for 5 minutes at 400g. The pellet was then plated without GM-CSF on glass coverslips on 24-well plates for phagocytosis assay.

### Syk inhibition

Highly-selective Syk inhibitor BAY-61-3606 (B9685, Sigma, St. Louis, MO) (Yamamoto et al., 2003), dissolved in 80% DMSO, was applied to N9 cells at different concentrations, ranging from 0.75 to 5µM for 2h, followed by the addition of aggregated human Aβ_42_ peptide at a concentration of 750nM (ab120301, Abcam, Cambridge, UK). Briefly, Aβ peptide was diluted in DMEM to a concentration of 7.5mM and incubated for 1h at 37°C to induce aggregation. Inhibition of Syk and downstream signaling molecules were assessed using western blotting and previously described.

### Protein extraction from cell-line and primary microglia

Cells were lysed in 0.1% SDS RIPA buffer (150mM NaCl, 5mM EDTA, 50mM Tris-base, 1% Triton, 0.5% Na-deoxycholate, 0.1% SDS in aqueous solution) containing protease inhibitor (1:100, P2714, Sigma, St. Louis, MO) and phosphate inhibitor (1:100, P5726, Sigma, St. Louis, MO) cocktails and incubated for 30min on ice, then centrifuged at 17,000g for 20min. The supernatant was separated, and total protein concentration was measured using the BCA method.

### Fluorescent-beads phagocytosis assay

N9 cells were deprived of FBS and incubated this Syk inhibitor BAY-61-3606 (5µM) for 3h. 1µm-fluorescent beads (L1030, Sigma, St. Louis, MO) were pre-coated with FBS for 1h at 37°C, then applied to cells at a dilution of 1:1000 for 1h at 37°C. Cells were then washed five times with PBS (containing calcium and magnesium) and trypsinized. Phagocytic FITC+ cell count was conducted using the BD-LSRFortessa cell analyzer (BD bioscience, East Rutherford, NJ).

### Aβ phagocytosis assay

Aβ_42_ peptide (ab120301, Abcam, Cambridge, UK) was diluted in DMEM and incubated for 3h at 37°C to induce aggregation. Aggregated Aβ was incubated with serum from anti-Aβ vaccinated mice, sham-vaccinated mice, or DMEM (500 ng/ml antibody, determined by ELISA) for 1h at 37°C with gentle agitation, to produce Ab-Ag complex. N9 and primary cells were treated with 5µM BAY-61-3606 for 3h for Syk inhibition and incubated with aggregated Aβ with or without the addition of serum for 4h. Next, cells were washed five times with PBS (containing calcium and magnesium). For immunofluorescence, cells were fixed with 4% PFA and permeabilized using 0.3% PBS-triton. Next, cells were stained with an anti-CD68, anti-Aβ1-14, and antibodies (Supplementary Table. 1), as previously described. Unbiased stereological counting of Aβ foci was performed using the Stereo-investigator software with a counting frame of 300×300µm. For FACS analysis, cells were fixed with 4% PFA, permeabilized using 0.3% PBS-triton, and stained for intracellular Aβ using an anti-Aβ1-14 antibody (Supplementary Table. 1). Aβ-containing cell count was conducted using the BD-LSRFortessa cell analyzer (BD bioscience, East Rutherford, NJ).

### Statistical analysis

The data presented as mean ± SEM were tested for significance in the unpaired *t*-test with equal variances, one-way ANOVA, repeated measures (RM) two-way ANOVA, two-sample Kolmogorov-Smirnov test, Pearson’s correlation coefficient, or the χ^2^ test for independence. *Post-hoc* tests were conducted using the Tukey or Bonferroni corrections. All error bars represent SEM were calculated as 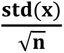 for numeric variables, and as 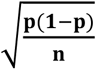 for binomial variables. Outliers were identified using the robust regression and outlier removal (ROUT) method with coefficient Q=1% (Motulsky & Brown, 2006). Significant results were marked according to conventional critical P values: *P<0.05, **P<0.01, ***P<0.001, ****P<0.0001.

## Data availability

All the data supporting the findings of this study as well as MATLAB codes are freely available upon request.

## Supporting information

Supplementary Text Results

## Acknowledgements

This study was conducted in the Paul Feder laboratory of Alzheimer’s disease research and was supported by the Clore Israel Foundation.

## Author contribution

TI and EO conceptualized and designed this study. TI, RN, and LBS performed the experiments. TI analyzed the data. RM provided methodological and biological insights and technical aid. AB developed the AβCoreS vaccine and provided mechanistic insights. TI and EO wrote the manuscript. All authors reviewed and approved the final version.

## Supplementary information titles and legends

### Supplementary text results

#### Supplementary figures

**Fig. S1. Maternally induced anti-Aβ antibodies cross the placenta and lactation into the circulation of 5xFAD fetuses and newborns.** (A) Color-coded heat maps of IgG isotypes in vaccinated dams prior to mating (left panel), isotypes in maternally vaccinated offspring at 1m of age, and following active vaccination at 2.5, 3.5, and 5m of age.

**Fig. S2. A combination of maternal and active vaccination rescues short-term memory abilities and normalizes exploratory behavior.** Supporting data for the main behavioral figure. (A) Speed and (B) distance traveled in the OF test. (C) Time spent in the open and close sections of the elevated zero-maze did not differ between groups, suggesting no differential effect of anxiety. (D) Distance and (E) speed in the EZM did not differ between groups. (F) Spontaneous alteration T-maze revealed no difference between groups. One-way ANOVA, two-way ANOVA.

**Fig. S3. Change in expression levels of hAPP does not account for cerebral Aβ levels following maternal and active vaccination.** (A) Transcript levels of cortical *hAPP* do not differ between groups. (B) Active vaccination reduces cerebral hAPP levels alongside with reducing Aβ. *P<0.05, two-way ANOVA.

**Fig. S4. Maternal vaccination but not active vaccination predicts reduced Aβ pathology at adulthood.** Scatter plot of insoluble Aβ_42_ levels and IgG isotypes following maternal (A-C) and active (D-F) vaccination. (A) Total IgG, (B) IgG2b, and (C) IgM at 1m of age negatively correlate with Aβ pathology at adulthood. Levels of (D) total IgG, (E) IgG2b, and (F) IgM at 5m of age poorly correlate with Aβ pathology, Pearson’s correlation.

**Fig. S5. Cerebral *FcγR* levels negatively correlate with Aβ pathology.** (A) Levels of *Fc**γ**R*s at 5m of age negatively correlate with cerebral Aβ pathology. (B-F) Scatter plot of insoluble Aβ_42_ levels and FcR expression levels at 5m of age present negative correlations. (B) *FcγRI*, (C) *FcγRIII,* (D) *FcγRIV,* (E) *FcRn*, and (F) *FcγRIIb*, Pearson’s correlation.

**Fig. S6. FcγRs are expressed in extra-follicular areas, the follicles and germinal centers in the spleen.** FcR Ab specificity was verified in spleen tissue slices prior to use on brain sections. (A) FcγRI, FcγRIII, FcγRIV, and FcRn were double-labeled with Iba1^+^ macrophages in the spleen. FcγRI is expressed on Iba1^+^ cells located at the surroundings of splenic follicles, and FcγRIIb, FcγRIII, and FcγRIV are co-localized with cells at extra-follicular areas, the follicles, and germinal centers.

**Fig. S7. FcγRI is expressed on microglial cells.** To assess FcγRI expression among CNS cells, double labeling was conducted with (A) Iba1^+^ (Microglia), NeuN (mature neurons) and GFAP (astrocytes), using the x10 objective. (B) FcγRI fluorescent signal intensity is higher in Iba1^+^ cells compared with both NeuN and GFAP^+^ cells. (C) Scatter plot and distributions of FcγRI expression on microglia, neurons, and astrocytes, showing normal distribution among microglia and right-skewed distributions for astrocytes and neurons (D) FcγRI is expressed within the somas of Iba1 ^+^cells. Images were taken using the x40 (left panels) and x63 (right panels) objectives. ****P<0.0001, one-way ANOVA.

**Fig. S8. FcγRI is upregulated among cortical microglia of maternally vaccinated 5xFAD mice.** (A) FcγRI expression was assessed using double-labeled immunofluorescence with Iba1^+^ microglia, using the x10 and x63 objectives for visualizing and quantification, respectively. (B) FcγRI signal was increased among microglia from maternally vaccinated mice independently of active vaccination. (C) Scatter plot of Iba1 and FcγRI signals reveal right-skewed distribution for FcγRI expression among unvaccinated and actively vaccinated mice, and normal distribution for maternally vaccinated mice. (D) Overlay and comparisons of FcγRI expression distribution. **P<0.01, ****P<0.0001, two-way ANOVA, corrected two-sample Kolmogorov-Smirnov test.

**Fig. S9. Maternal vaccination activates FcR-mediated phagocytosis via activation of AKT, ERK, and Cofilin actin-cytoskeleton regulation pathways.** Levels of Syk and downstream signaling molecules from the Fc-mediated phagocytosis pathway were measured using immunoblotting following maternal vaccination and active vaccination at 5m of age (see main Fig. 5A). (A) Maternal and active vaccination activates FcγR-mediated phagocytosis pathways through actin cytoskeleton regulation.

**Fig. S10. Maternal vaccination increases Syk activation in hippocampal microglia.** (A) Syk activation among microglial cells was assessed using double labeling of pSyk and CD68 positive microglia, using the x10 and x63 objectives for visualizing and quantification, respectively. (B) pSyk signal was higher in maternally and actively immunized mice compared with unvaccinated controls. *P<0.05, ****P<0.0001, two-way ANOVA.

**Fig. S11. Dose-dependent Syk inhibition by BAY-61-3606 in the N9 microglial cell line.** BAY-61-3606 was applied to N9 cells at different concentrations, ranging from 0.75 to 5µM for 2h, followed by the addition of aggregated human Aβ_42_ peptide at a concentration of 750nM. (A-C) Western blotting of phospho- and total Syk and downstream signaling molecules from the FcR mediated phagocytosis pathway: AKT, ERK, and Cofilin.

#### Supplementary tables

Supplementary Table 1. sELISA, IF and WB primary antibodies

Supplementary Table 2. RT-PCR primers

Supplementary materials and methods

## Reference

1. ACOG. (2018). ACOG Committee Opinion No. 741: Maternal Immunization. Obstet Gynecol, 131(6), e214–e217. doi: 10.1097/AOG.0000000000002662

2. Alhajraf, F., Ness, D., Hye, A., & Strydom, A. (2019). Plasma amyloid and tau as dementia biomarkers in Down syndrome: systematic review and meta-analyses. Dev Neurobiol. doi: 10.1002/dneu.22715

3. Atwood, C. S., Moir, R. D., Huang, X., Scarpa, R. C., Bacarra, N. M., Romano, D. M., … Bush, A. I. (1998). Dramatic aggregation of Alzheimer abeta by Cu(II) is induced by conditions representing physiological acidosis. J Biol Chem, 273(21), 12817–12826. doi: 10.1074/jbc.273.21.12817

4. Barone, E., Head, E., Butterfield, D. A., & Perluigi, M. (2017). HNE-modified proteins in Down syndrome: Involvement in development of Alzheimer disease neuropathology. Free Radic Biol Med, 111, 262–269. doi: 10.1016/j.freeradbiomed.2016.10.508

5. Bruhns, P., & Jonsson, F. (2015). Mouse and human FcR effector functions. Immunol Rev, 268(1), 25–51. doi: 10.1111/imr.12350

6. Burger, P. C., & Vogel, F. S. (1973). The development of the pathologic changes of Alzheimer’s disease and senile dementia in patients with Down’s syndrome. Am J Pathol, 73(2), 457–476.

7. Butterfield, D. A., & Perluigi, M. (2018). Down syndrome: From development to adult life to Alzheimer disease. Free Radic Biol Med, 114, 1–2. doi: 10.1016/j.freeradbiomed.2017.10.374

8. Carfi, A., Antocicco, M., Brandi, V., Cipriani, C., Fiore, F., Mascia, D., … Onder, G. (2014). Characteristics of adults with down syndrome: prevalence of age-related conditions. Front Med (Lausanne), 1, 51. doi: 10.3389/fmed.2014.00051

9. Chan, S. L., Furukawa, K., & Mattson, M. P. (2002). Presenilins and APP in neuritic and synaptic plasticity: implications for the pathogenesis of Alzheimer’s disease. Neuromolecular Med, 2(2), 167–196. doi: 10.1385/NMM:2:2:167

10. Chistiakov, D. A., Killingsworth, M. C., Myasoedova, V. A., Orekhov, A. N., & Bobryshev, Y. V. (2017). CD68/macrosialin: not just a histochemical marker. Lab Invest, 97(1), 4–13. doi: 10.1038/labinvest.2016.116

11. Choong, X. Y., Tosh, J. L., Pulford, L. J., & Fisher, E. M. (2015). Dissecting Alzheimer disease in Down syndrome using mouse models. Front Behav Neurosci, 9, 268. doi: 10.3389/fnbeh.2015.00268

12. Corradin, S. B., Mauel, J., Donini, S. D., Quattrocchi, E., & Ricciardi-Castagnoli, P. (1993). Inducible nitric oxide synthase activity of cloned murine microglial cells. Glia, 7(3), 255–262. doi: 10.1002/glia.440070309

13. Deacon, R. M. J., & Rawlins, J. N. P. (2006). T-maze alternation in the rodent. Nat. Protocols, 1(1), 7–12.

14. Fu, R., Shen, Q., Xu, P., Luo, J. J., & Tang, Y. (2014). Phagocytosis of microglia in the central nervous system diseases. Mol Neurobiol, 49(3), 1422–1434. doi: 10.1007/s12035-013-8620-6

15. Geylis, V., & Steinitz, M. (2006). Immunotherapy of Alzheimer’s disease (AD): from murine models to anti-amyloid beta (Abeta) human monoclonal antibodies. Autoimmun Rev, 5(1), 33–39. doi: 10.1016/j.autrev.2005.06.007

16. Gribble, S. M., Wiseman, F. K., Clayton, S., Prigmore, E., Langley, E., Yang, F., … Carter, N. P. (2013). Massively parallel sequencing reveals the complex structure of an irradiated human chromosome on a mouse background in the Tc1 model of Down syndrome. PLoS One, 8(4), e60482. doi: 10.1371/journal.pone.0060482

17. Gupta, M., Dhanasekaran, A. R., & Gardiner, K. J. (2016). Mouse models of Down syndrome: gene content and consequences. Mamm Genome, 27(11-12), 538–555. doi: 10.1007/s00335-016-9661-8

18. Heneka, M. T. (2019). Microglia take centre stage in neurodegenerative disease. Nat Rev Immunol, 19(2), 79–80. doi: 10.1038/s41577-018-0112-5

19. Illouz, T., Madar, R., Biragyn, A., & Okun, E. (2019). Restoring microglial and astroglial homeostasis using DNA immunization in a Down Syndrome mouse model. Brain Behav Immun, 75, 163–180. doi: 10.1016/j.bbi.2018.10.004

20. Illouz, T., Madar, R., Griffioen, K., & Okun, E. (2017). A protocol for quantitative analysis of murine and human amyloid-beta1-40 and 1-42. J Neurosci Methods, 291, 28–35. doi: 10.1016/j.jneumeth.2017.07.022

21. Jankowsky, J. L., Younkin, L. H., Gonzales, V., Fadale, D. J., Slunt, H. H., Lester, H. A., … Borchelt, D. R. (2007). Rodent A beta modulates the solubility and distribution of amyloid deposits in transgenic mice. J Biol Chem, 282(31), 22707–22720. doi: 10.1074/jbc.M611050200

22. Jarrett, J. T., & Lansbury, P. T., Jr. (1993). Seeding “one-dimensional crystallization” of amyloid: a pathogenic mechanism in Alzheimer’s disease and scrapie? Cell, 73(6), 1055–1058. doi: 10.1016/0092-8674(93)90635-4

23. Jaumouille, V., Farkash, Y., Jaqaman, K., Das, R., Lowell, C. A., & Grinstein, S. (2014). Actin cytoskeleton reorganization by Syk regulates Fcgamma receptor responsiveness by increasing its lateral mobility and clustering. Dev Cell, 29(5), 534–546. doi: 10.1016/j.devcel.2014.04.031

24. Kanellos, G., & Frame, M. C. (2016). Cellular functions of the ADF/cofilin family at a glance. J Cell Sci, 129(17), 3211–3218. doi: 10.1242/jcs.187849

25. Keren-Shaul, H., Spinrad, A., Weiner, A., Matcovitch-Natan, O., Dvir-Szternfeld, R., Ulland, T. K., … Amit, I. (2017). A Unique Microglia Type Associated with Restricting Development of Alzheimer’s Disease. Cell, 169(7), 1276–1290 e1217. doi: 10.1016/j.cell.2017.05.018

26. Kiefer, F., Brumell, J., Al-Alawi, N., Latour, S., Cheng, A., Veillette, A., … Pawson, T. (1998). The Syk protein tyrosine kinase is essential for Fcgamma receptor signaling in macrophages and neutrophils. Mol Cell Biol, 18(7), 4209–4220. doi: 10.1128/mcb.18.7.4209

27. Korbel, J. O., Tirosh-Wagner, T., Urban, A. E., Chen, X. N., Kasowski, M., Dai, L., … Korenberg, J. R. (2009). The genetic architecture of Down syndrome phenotypes revealed by high-resolution analysis of human segmental trisomies. Proc Natl Acad Sci U S A, 106(29), 12031–12036. doi: 10.1073/pnas.0813248106

28. Krasemann, S., Madore, C., Cialic, R., Baufeld, C., Calcagno, N., El Fatimy, R., … Butovsky, O. (2017). The TREM2-APOE Pathway Drives the Transcriptional Phenotype of Dysfunctional Microglia in Neurodegenerative Diseases. Immunity, 47(3), 566–581 e569. doi: 10.1016/j.immuni.2017.08.008

29. Latvala, S., Jacobsen, B., Otteneder, M. B., Herrmann, A., & Kronenberg, S. (2017). Distribution of FcRn Across Species and Tissues. J Histochem Cytochem, 65(6), 321–333. doi: 10.1369/0022155417705095

30. Leger, M., Quiedeville, A., Bouet, V., Haelewyn, B., Boulouard, M., Schumann-Bard, P., & Freret, T. (2013). Object recognition test in mice. Nat Protoc, 8(12), 2531–2537. doi: 10.1038/nprot.2013.155

31. Lueptow, L. M. (2017). Novel Object Recognition Test for the Investigation of Learning and Memory in Mice. J Vis Exp(126). doi: 10.3791/55718

32. Miller, G. (2012). Alzheimer’s research. Stopping Alzheimer’s before it starts. Science, 337(6096), 790–792. doi: 10.1126/science.337.6096.790

33. Morris, J. C., Aisen, P. S., Bateman, R. J., Benzinger, T. L., Cairns, N. J., Fagan, A. M., … Buckles, V. D. (2012). Developing an international network for Alzheimer research: The Dominantly Inherited Alzheimer Network. Clin Investig (Lond), 2(10), 975–984. doi: 10.4155/cli.12.93

34. Motulsky, H. J., & Brown, R. E. (2006). Detecting outliers when fitting data with nonlinear regression - a new method based on robust nonlinear regression and the false discovery rate. BMC Bioinformatics, 7, 123. doi: 10.1186/1471-2105-7-123

35. Moussaud, S., & Draheim, H. J. (2010). A new method to isolate microglia from adult mice and culture them for an extended period of time. J Neurosci Methods, 187(2), 243–253. doi: 10.1016/j.jneumeth.2010.01.017

36. Munoz, F. M., & Jamieson, D. J. (2019). Maternal Immunization. Obstet Gynecol, 133(4), 739–753. doi: 10.1097/AOG.0000000000003161

37. Nilaratanakul, V., Chen, J., Tran, O., Baxter, V. K., Troisi, E. M., Yeh, J. X., & Griffin, D. E. (2018). Germ Line IgM Is Sufficient, but Not Required, for Antibody-Mediated Alphavirus Clearance from the Central Nervous System. J Virol, 92(7). doi: 10.1128/JVI.02081-17

38. Oakley, H., Cole, S. L., Logan, S., Maus, E., Shao, P., Craft, J., … Vassar, R. (2006). Intraneuronal beta-amyloid aggregates, neurodegeneration, and neuron loss in transgenic mice with five familial Alzheimer’s disease mutations: potential factors in amyloid plaque formation. J Neurosci, 26(40), 10129–10140. doi: 10.1523/JNEUROSCI.1202-06.2006

39. Olkhanud, P. B., Mughal, M., Ayukawa, K., Malchinkhuu, E., Bodogai, M., Feldman, N., … Biragyn, A. (2012). DNA immunization with HBsAg-based particles expressing a B cell epitope of amyloid beta-peptide attenuates disease progression and prolongs survival in a mouse model of Alzheimer’s disease. Vaccine, 30(9), 1650–1658. doi: 10.1016/j.vaccine.2011.12.136

40. Presson, A. P., Partyka, G., Jensen, K. M., Devine, O. J., Rasmussen, S. A., McCabe, L. L., & McCabe, E. R. (2013). Current estimate of Down Syndrome population prevalence in the United States. J Pediatr, 163(4), 1163–1168. doi: 10.1016/j.jpeds.2013.06.013

41. Pyrski, M., Rugowska, A., Wierzbinski, K. R., Kasprzyk, A., Bogusiewicz, M., Bociag, P., … Pniewski, T. (2017). HBcAg produced in transgenic tobacco triggers Th1 and Th2 response when intramuscularly delivered. Vaccine, 35(42), 5714–5721. doi: 10.1016/j.vaccine.2017.07.082

42. Raha-Chowdhury, R., Henderson, J. W., Raha, A. A., Stott, S. R. W., Vuono, R., Foscarin, S., … Zaman, S. H. (2018). Erythromyeloid-Derived TREM2: A Major Determinant of Alzheimer’s Disease Pathology in Down Syndrome. J Alzheimers Dis, 61(3), 1143–1162. doi: 10.3233/JAD-170814

43. Reiman, E. M., Langbaum, J. B., Fleisher, A. S., Caselli, R. J., Chen, K., Ayutyanont, N., … Tariot, P. N. (2011). Alzheimer’s Prevention Initiative: a plan to accelerate the evaluation of presymptomatic treatments. J Alzheimers Dis, 26 *Suppl 3*, 321–329. doi: 10.3233/JAD-2011-0059

44. Rosenberg, R. N., Fu, M., & Lambracht-Washington, D. (2018). Intradermal active full-length DNA Abeta42 immunization via electroporation leads to high anti-Abeta antibody levels in wild-type mice. J Neuroimmunol, 322, 15–25. doi: 10.1016/j.jneuroim.2018.05.017

45. Rueda, N., Florez, J., & Martinez-Cue, C. (2012). Mouse models of Down syndrome as a tool to unravel the causes of mental disabilities. Neural Plast, 2012, 584071. doi: 10.1155/2012/584071

46. Seibenhener, M. L., & Wooten, M. C. (2015). Use of the Open Field Maze to measure locomotor and anxiety-like behavior in mice. J Vis Exp(96), e52434. doi: 10.3791/52434

47. Shepherd, J. K., Grewal, S. S., Fletcher, A., Bill, D. J., & Dourish, C. T. (1994). Behavioural and pharmacological characterisation of the elevated “zero-maze” as an animal model of anxiety. Psychopharmacology (Berl), 116(1), 56–64. doi: 10.1007/bf02244871

48. Song, X., Tanaka, S., Cox, D., & Lee, S. C. (2004). Fcgamma receptor signaling in primary human microglia: differential roles of PI-3K and Ras/ERK MAPK pathways in phagocytosis and chemokine induction. J Leukoc Biol, 75(6), 1147–1155. doi: 10.1189/jlb.0403128

49. Stangel, M., & Compston, A. (2001). Polyclonal immunoglobulins (IVIg) modulate nitric oxide production and microglial functions in vitro via Fc receptors. J Neuroimmunol, 112(1-2), 63–71. doi: 10.1016/s0165-5728(00)00412-4

50. Strohmeyer, R., Kovelowski, C. J., Mastroeni, D., Leonard, B., Grover, A., & Rogers, J. (2005). Microglial responses to amyloid beta peptide opsonization and indomethacin treatment. J Neuroinflammation, 2, 18. doi: 10.1186/1742-2094-2-18

51. Takai, T. (2005). Fc receptors and their role in immune regulation and autoimmunity. J Clin Immunol, 25(1), 1–18. doi: 10.1007/s10875-005-0353-8

52. Tanaka, K., Takeda, S., Mitsuoka, K., Oda, T., Kimura-Sakiyama, C., Maeda, Y., & Narita, A. (2018). Structural basis for cofilin binding and actin filament disassembly. Nat Commun, 9(1), 1860. doi: 10.1038/s41467-018-04290-w

53. Tejera, D., & Heneka, M. T. (2019). Microglia in Neurodegenerative Disorders. Methods Mol Biol, 2034, 57–67. doi: 10.1007/978-1-4939-9658-2_5

54. Tramutola, A., Lanzillotta, C., Barone, E., Arena, A., Zuliani, I., Mosca, L., … Di Domenico, F. (2018). Intranasal rapamycin ameliorates Alzheimer-like cognitive decline in a mouse model of Down syndrome. Transl Neurodegener, 7, 28. doi: 10.1186/s40035-018-0133-9

55. Wiseman, F. K., Al-Janabi, T., Hardy, J., Karmiloff-Smith, A., Nizetic, D., Tybulewicz, V. L., … Strydom, A. (2015). A genetic cause of Alzheimer disease: mechanistic insights from Down syndrome. Nat Rev Neurosci, 16(9), 564–574. doi: 10.1038/nrn3983

56. Wiseman, F. K., Pulford, L. J., Barkus, C., Liao, F., Portelius, E., Webb, R., … LonDown, S. C. (2018). Trisomy of human chromosome 21 enhances amyloid-beta deposition independently of an extra copy of APP. Brain, 141(8), 2457–2474. doi: 10.1093/brain/awy159

57. Yamamoto, N., Takeshita, K., Shichijo, M., Kokubo, T., Sato, M., Nakashima, K., … Bacon, K. B. (2003). The orally available spleen tyrosine kinase inhibitor 2-[7-(3,4-dimethoxyphenyl)-imidazo[1,2-c]pyrimidin-5-ylamino]nicotinamide dihydrochloride (BAY 61-3606) blocks antigen-induced airway inflammation in rodents. J Pharmacol Exp Ther, 306(3), 1174–1181. doi: 10.1124/jpet.103.052316

