## Supplementary Text Results for "Maternal antibodies facilitate Amyloid-β clearance by activating Fc-receptor-Syk-mediated phagocytosis"

**Effector *FcR* transcript levels negatively correlate with Aβ load in the brain.** We assessed whether upregulation of FcRs was correlated with a reduction in Aβ load. *FcγRI*, *FcγRIII*, *FcγRIV*, *FcRn* and *FcγRIIb* mRNA levels negatively correlated with cortical SDS-soluble $Aβ_{40}$ levels (*r*=-0.64, P=0.001, *r*=-0.63, P=0.001, *r*=-0.43, P=0.02, *r*=-0.59, P=0.002, *r*=-0.56, P=0.004, respectively, Fig. S5A, left panel). Levels of TBST-soluble $Aβ_{42}$ negatively correlated with mRNA levels of *FcγRI* and *FcγRIII* (r=-0.43, P=0.02, r=-0.4, P=0.03, respectively, Fig. S5A, right panel) but not with *FcγRIV*, *FcRn*, or *FcγRIIb*. Additionally, *FcγRI*, *FcγRIII*, *FcγRIV*, *FcRn* and *FcγRIIb* mRNA levels negatively correlated with cortical SDS-soluble $Aβ_{42}$ levels (*r*=-0.57, P=0.004, *r*=-0.53, P=0.007, *r*=-0.51, P=0.009, *r*=-0.6, P=0.002, *r*=0.44, P=0.03, respectively, Fig. S5A, right panel). Importantly, *FcγRI*, *FcγRIII*, and *FcRn* were negatively correlated with insoluble $Aβ_{42}$ (*r*=-0.42, P=0.03, *r*=-0.4, P=0.03, *r*=-0.42, P=0.02, Fig. S5A, right panel, S4B, C, E). *FcγRIV* and *FcγRIIb* did not correlate with insoluble $Aβ_{42}$ (Fig. S5A, right panel, S4D, F).

**Maternal vaccination elevates FcγRI on microglial cells.** Ab specificity for these receptors was verified on mouse spleen sections (Fig. S6). Of these receptors, FcγRI was the only receptor found to be expressed in the brain (Fig. S7). Co-immunostaining with the cell-specific Iba1, NeuN, and GFAP markers revealed that FcγRI was exclusively expressed on $Iba^+$ microglia, but not $Neun^+$ neurons or $GFAP^+$ astrocytes (33.01±7.49, 2.7±1.23, 1.34±2.3 a.u (arbitrary units), P<0.0001, Fig. S7A-B). The distribution of cellular expression on $Iba^+$, but not on $NeuN^+$ and $GFAP^+$, appears to be close to normal (mean=33.01, median=32.32, Fig. S7C, D), with no significant difference from a simulated normal distribution with the same mean and standard deviation (P=0.26, Fig S7C). FcγRI was thus expressed mainly on the vast majority of microglial cells, with its expression intensity normally distributed (Fig. S7C, D).

**FcγRI is elevated in cortical microglia of maternally vaccinated mice.** As seen in the hippocampus, cortiacal FcγRI levels were elevated in maternally vaccinated mice M+/A- and M+/A+ compared with both unvaccinated M-/A- and actively vaccinated mice M-/A+ (19.98±2.29, 18.35±2.09, 9.57±2.02, 13.78±1.59 a.u, $P<0.01$, Fig. S8A-B). FcγRI distribution is normal among all vaccinated groups ($P=0.09$, $P=0.2$, $P=0.14$, compared to a simulated normal distribution, respectively, Fig. S8A, C), while this distribution in unvaccinated mice appears to be skewed ($P<0.05$, Fig. S8A, C). Active vaccination alone seems to elevate levels of microglial FcγRI compared with unvaccinated mice. This elevation did not reach significance in two-way ANOVA, although the distributions of the two groups did significantly differ ($P<0.0001$, Fig. S8D).
