## Supplementary Figures for "Maternal antibodies facilitate Amyloid-β clearance by activating Fc-receptor-Syk-mediated phagocytosis"

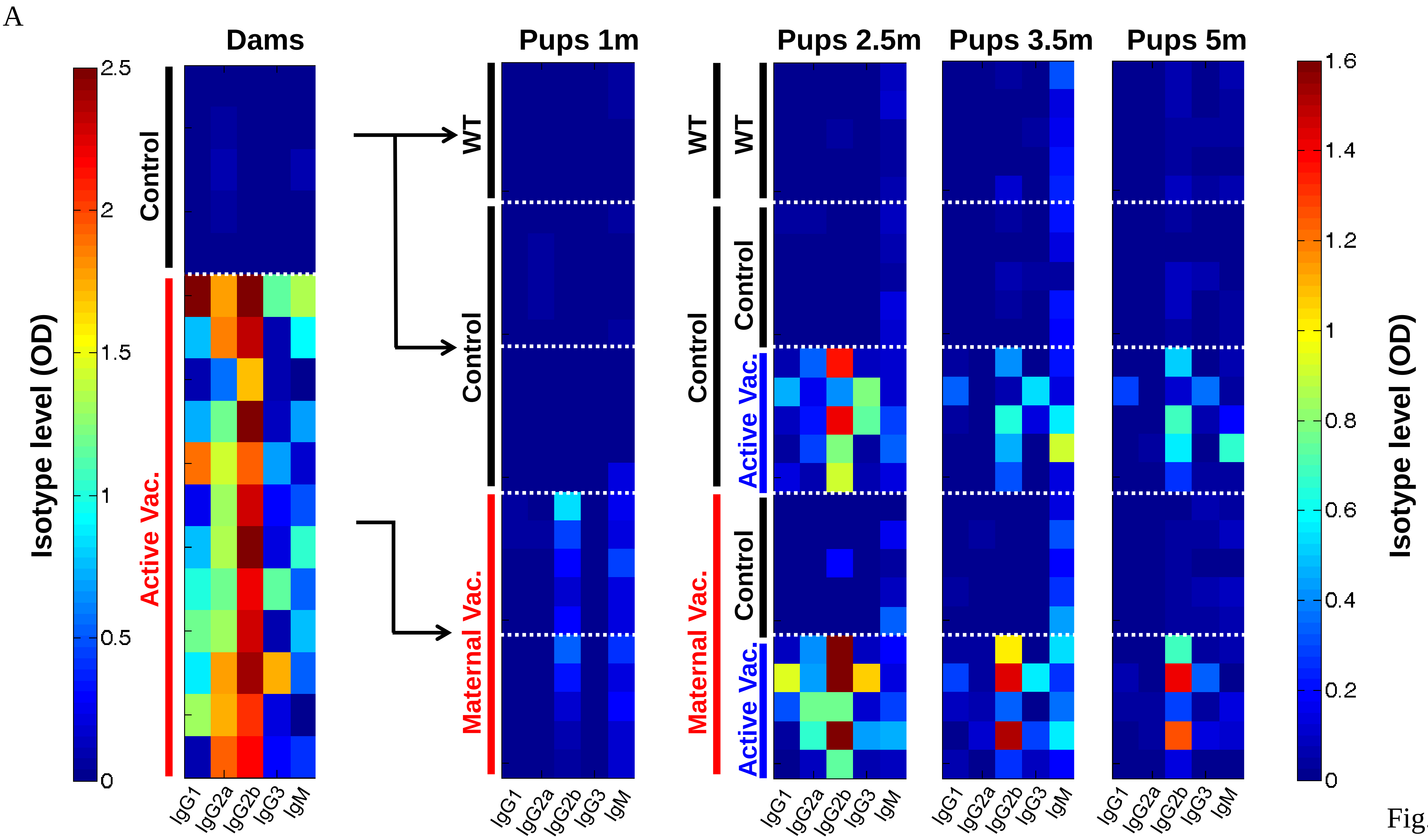
A
Dams
Pups 1m
Pups 2.5m
Pups 3.5m
Pups 5m
Isotype level (OD)
2.5
2
1.5
1
0.5
0
Control
Active Vac.
WT
Control
Maternal Vac.
Active Vac.
IgG1
IgG2a
IgG2b
IgG3
IgM
1.6
1.4
1.2
1
0.8
0.6
0.4
0.2
0
Isotype level (OD)

Fig. S1

**Fig. S1. Maternally induced anti-Aβ antibodies cross the placenta and lactation into the circulation of 5xFAD fetuses and newborns.** (A) Color-coded heat maps of IgG isotypes in vaccinated dams prior to mating (left panel), isotypes in maternally vaccinated offspring at 1m of age, and following active vaccination at 2.5, 3.5, and 5m of age.

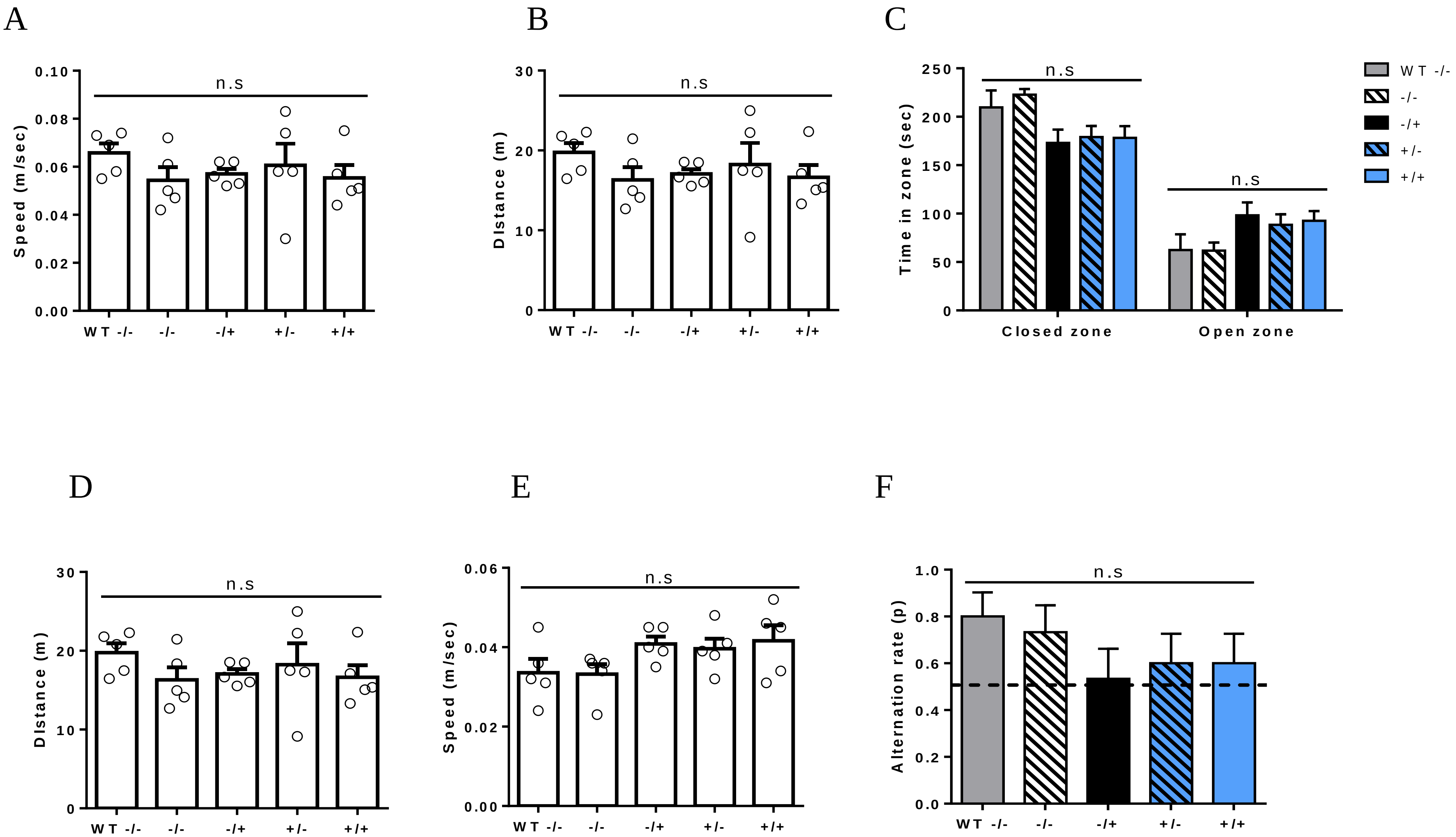
A
B
C
D
E
F
n.s
Speed (m/sec)
0.10
0.08
0.06
0.04
0.02
0.00
WT -/-
-/-
-/+
+/-
+/+
Distance (m)
30
20
10
0
Time in zone (sec)
250
200
150
100
50
Closed zone
Open zone
Alternation rate (p)
1.0
0.8
0.6
0.4
0.2
0.0

Fig. S2

**Fig. S2. A combination of maternal and active vaccination rescues short-term memory abilities and normalizes exploratory behavior.** Supporting data for the main behavioral figure. (A) Speed and (B) distance traveled in the OF test. (C) Time spent in the open and close sections of the elevated zero-maze did not differ between groups, suggesting no differential effect of anxiety. (D) Distance and (E) speed in the EZM did not differ between groups. (F) Spontaneous alteration T-maze revealed no difference between groups. One-way ANOVA, two-way ANOVA.

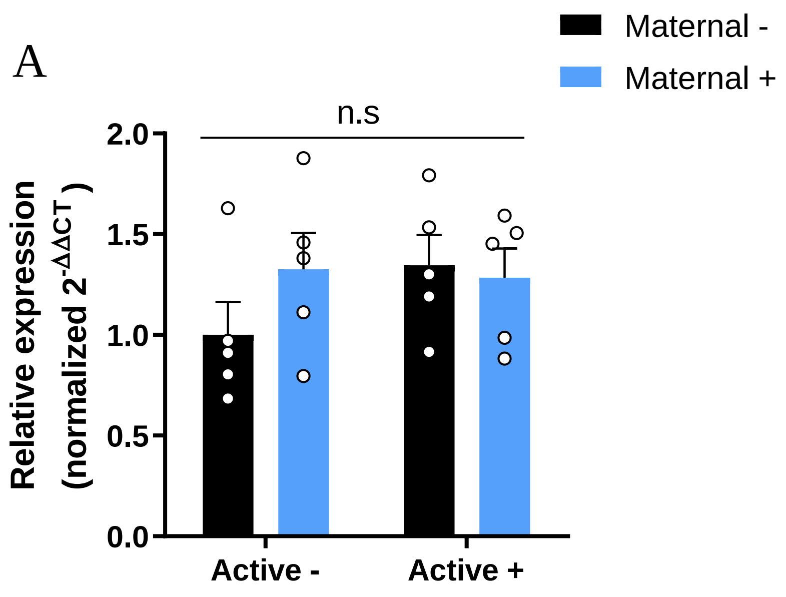
A
Maternal -
Maternal +
n.s
Relative expression (normalized 2-ΔΔCT)
2.0
1.5
1.0
0.5
0.0
Active -
Active +

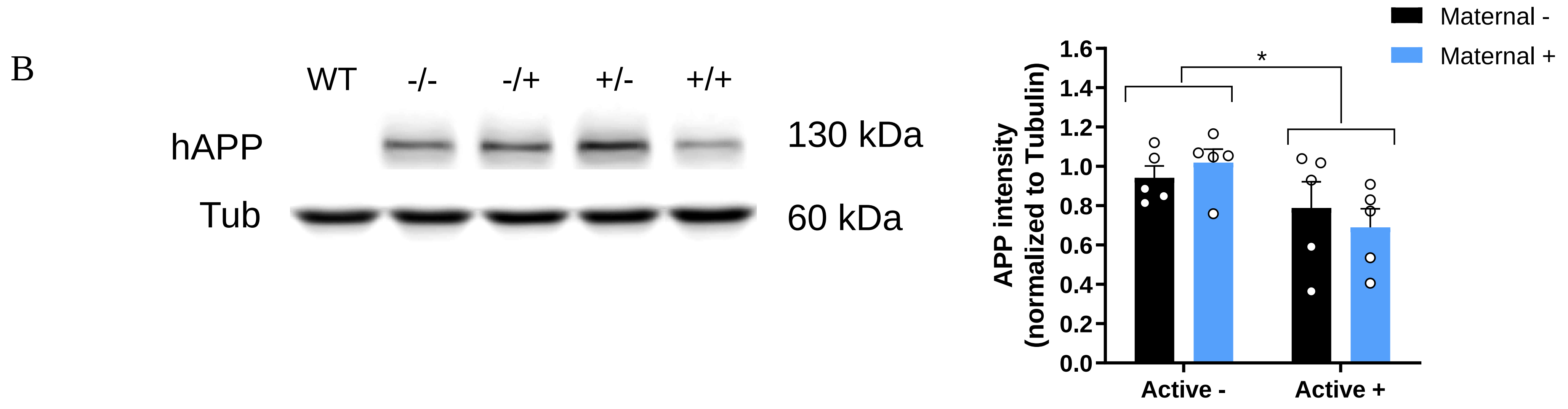
B
WT
-/-
-/+
+/-
+/+
hAPP
Tub
130 kDa
60 kDa
Maternal -
Maternal +
*
APP intensity (normalized to Tubulin)
1.6
1.4
1.2
1.0
0.8
0.6
0.4
0.2
0.0
Active -
Active +

Fig. S3

**Fig. S3. Change in expression levels of hAPP does not account for cerebral Aβ levels following maternal and active vaccination.** (A) Transcript levels of cortical *hAPP* do not differ between groups. (B) Active vaccination reduces cerebral hAPP levels alongside with reducing Aβ. *$P<0.05$, two-way ANOVA.

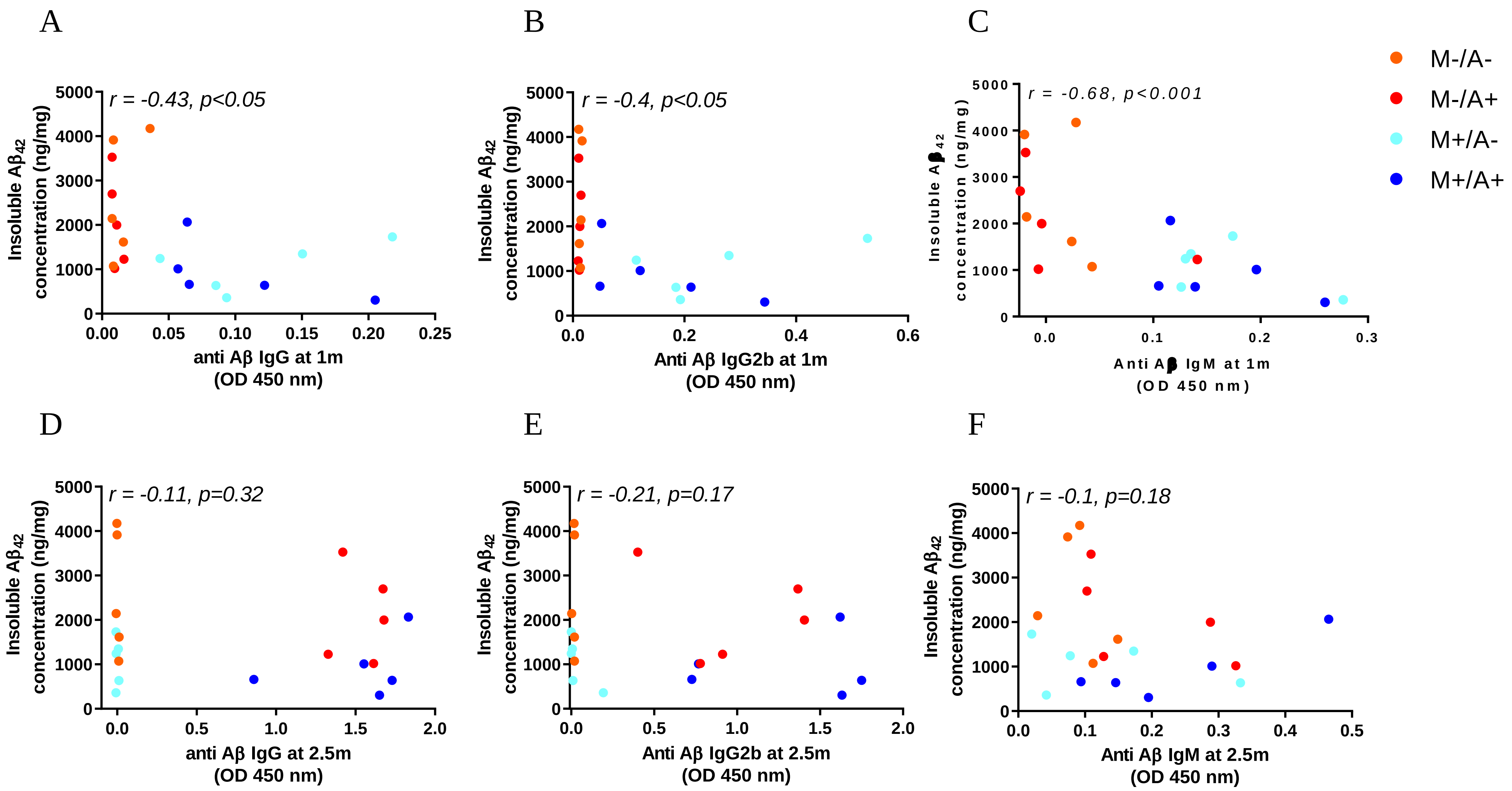
A
r = -0.43, p<0.05
Insoluble Aβ42 concentration (ng/mg)
anti Aβ IgG at 1m (OD 450 nm)
B
r = -0.4, p<0.05
Insoluble Aβ42 concentration (ng/mg)
Anti Aβ IgG2b at 1m (OD 450 nm)
C
r = -0.68, p<0.001
Insoluble Aβ42 concentration (ng/mg)
Anti Aβ IgM at 1m (OD 450 nm)
M-/A-
M-/A+
M+/A-
M+/A+
D
r = -0.11, p=0.32
Insoluble Aβ42 concentration (ng/mg)
anti Aβ IgG at 2.5m (OD 450 nm)
E
r = -0.21, p=0.17
Insoluble Aβ42 concentration (ng/mg)
Anti Aβ IgG2b at 2.5m (OD 450 nm)
F
r = -0.1, p=0.18
Insoluble Aβ42 concentration (ng/mg)
Anti Aβ IgM at 2.5m (OD 450 nm)

Fig. S4

**Fig. S4. Maternal vaccination but not active vaccination predicts reduced Aβ pathology at adulthood.** Scatter plot of insoluble $A\beta_{42}$ levels and IgG isotypes following maternal (A-C) and active (D-F) vaccination. (A) Total IgG, (B) IgG2b, and (C) IgM at 1m of age negatively correlate with Aβ pathology at adulthood. Levels of (D) total IgG, (E) IgG2b, and (F) IgM at 5m of age poorly correlate with Aβ pathology, Pearson's correlation.

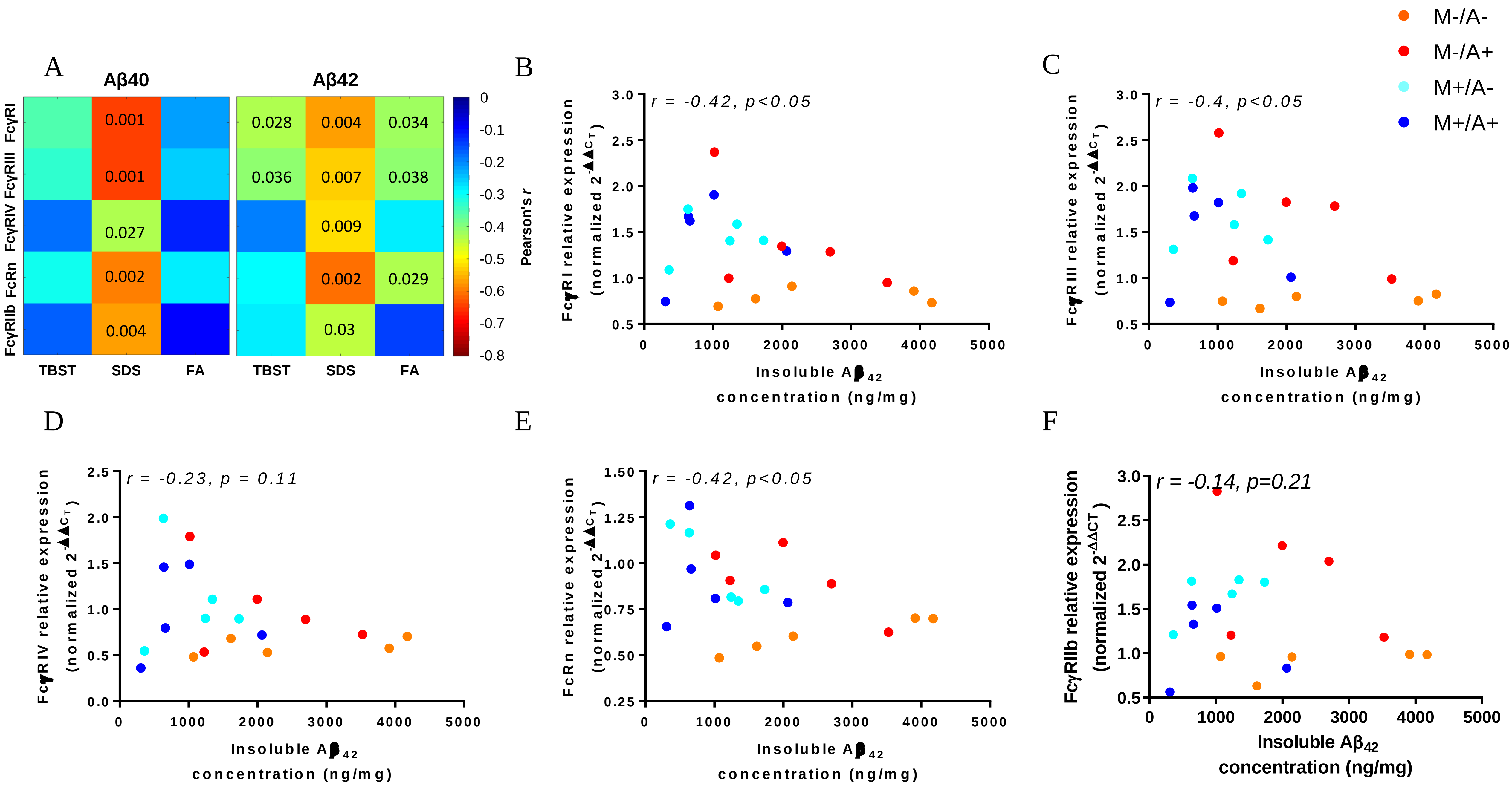

A
Aβ40
Aβ42
FcγRI
FcγRIII
FcγRIV
FcRn
FcγRIIb
0.001
0.001
0.027
0.002
0.004
0.028
0.004
0.034
0.036
0.007
0.038
0.009
0.002
0.029
0.03
TBST
SDS
FA
Pearson's r
B
r = -0.42, p<0.05
FcγRI relative expression (normalized 2-ΔΔCT)
Insoluble Aβ42 concentration (ng/mg)
C
r = -0.4, p<0.05
FcγRIII relative expression (normalized 2-ΔΔCT)
M-/A-
M-/A+
M+/A-
M+/A+
D
r = -0.23, p = 0.11
FcγRIV relative expression (normalized 2-ΔΔCT)
E
r = -0.42, p<0.05
FcRn relative expression (normalized 2-ΔΔCT)
F
r = -0.14, p=0.21
FcγRIIb relative expression (normalized 2-ΔΔCT)


Fig. S5

**Fig. S5. Cerebral *FcγR* levels negatively correlate with Aβ pathology.** (A) Levels of *FcγR*s at 5m of age negatively correlate with cerebral Aβ pathology. (B-F) Scatter plot of insoluble $A\beta_{42}$ levels and FcR expression levels at 5m of age present negative correlations. (B) *FcγRI*, (C) *FcγRIII*, (D) *FcγRIV*, (E) *FcRn*, and (F) *FcγRIIb*, Pearson's correlation.

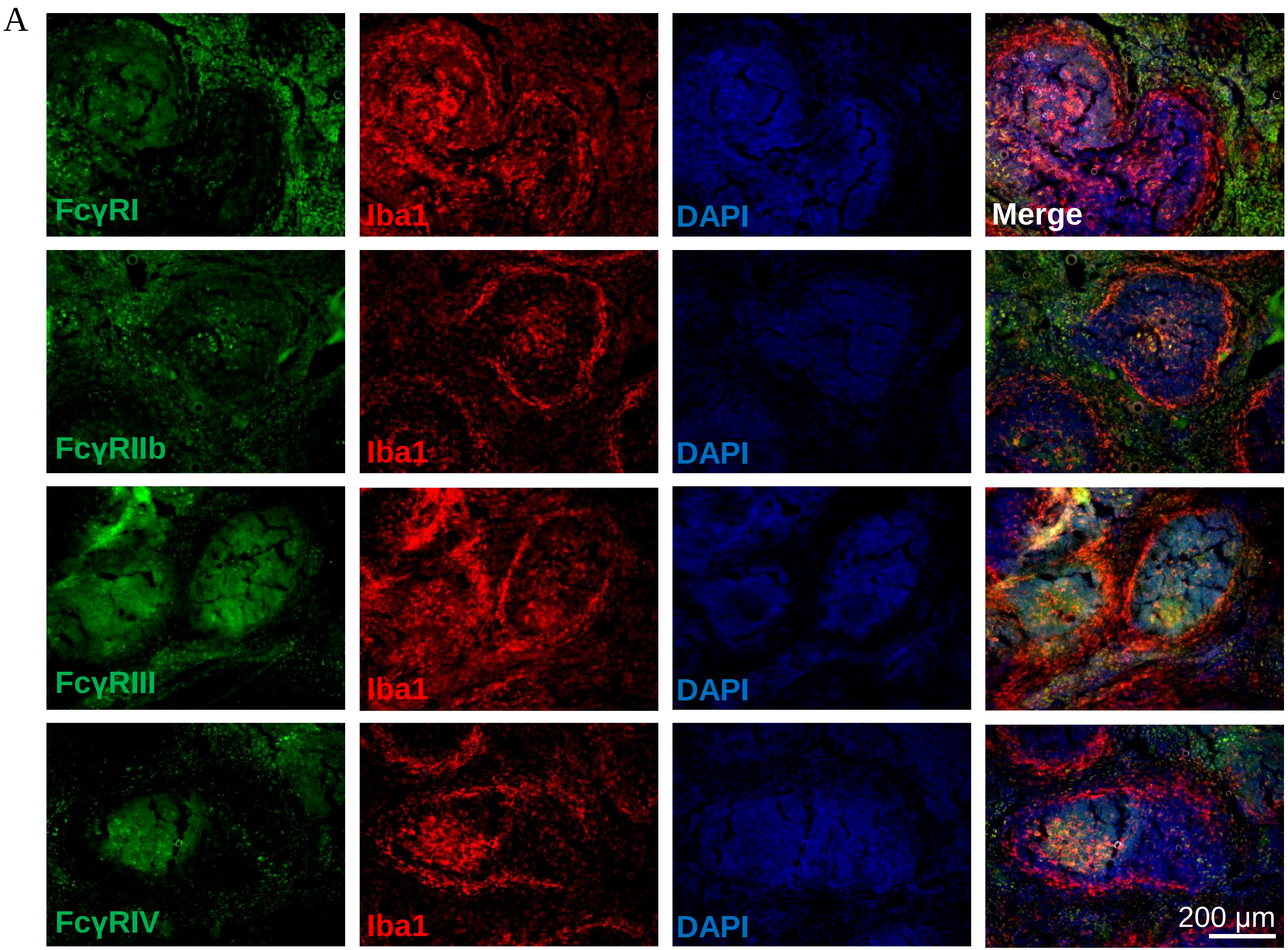
A
FcγRI
Iba1
DAPI
Merge
FcγRIIb
Iba1
DAPI
FcγRIII
Iba1
DAPI
FcγRIV
Iba1
DAPI
200 μm

Fig. S6

**Fig. S6. FcγRs are expressed in extra-follicular areas, the follicles and germinal centers in the spleen.** FcR Ab specificity was verified in spleen tissue slices prior to use on brain sections. (A) FcγRI, FcγRIII, FcγRIV, and FcRn were double-labeled with $Iba1^{+}$ macrophages in the spleen. FcγRI is expressed on $Iba1^{+}$ cells located at the surroundings of splenic follicles, and FcγRIIb, FcγRIII, and FcγRIV are co-localized with cells at extra-follicular areas, the follicles, and germinal centers.

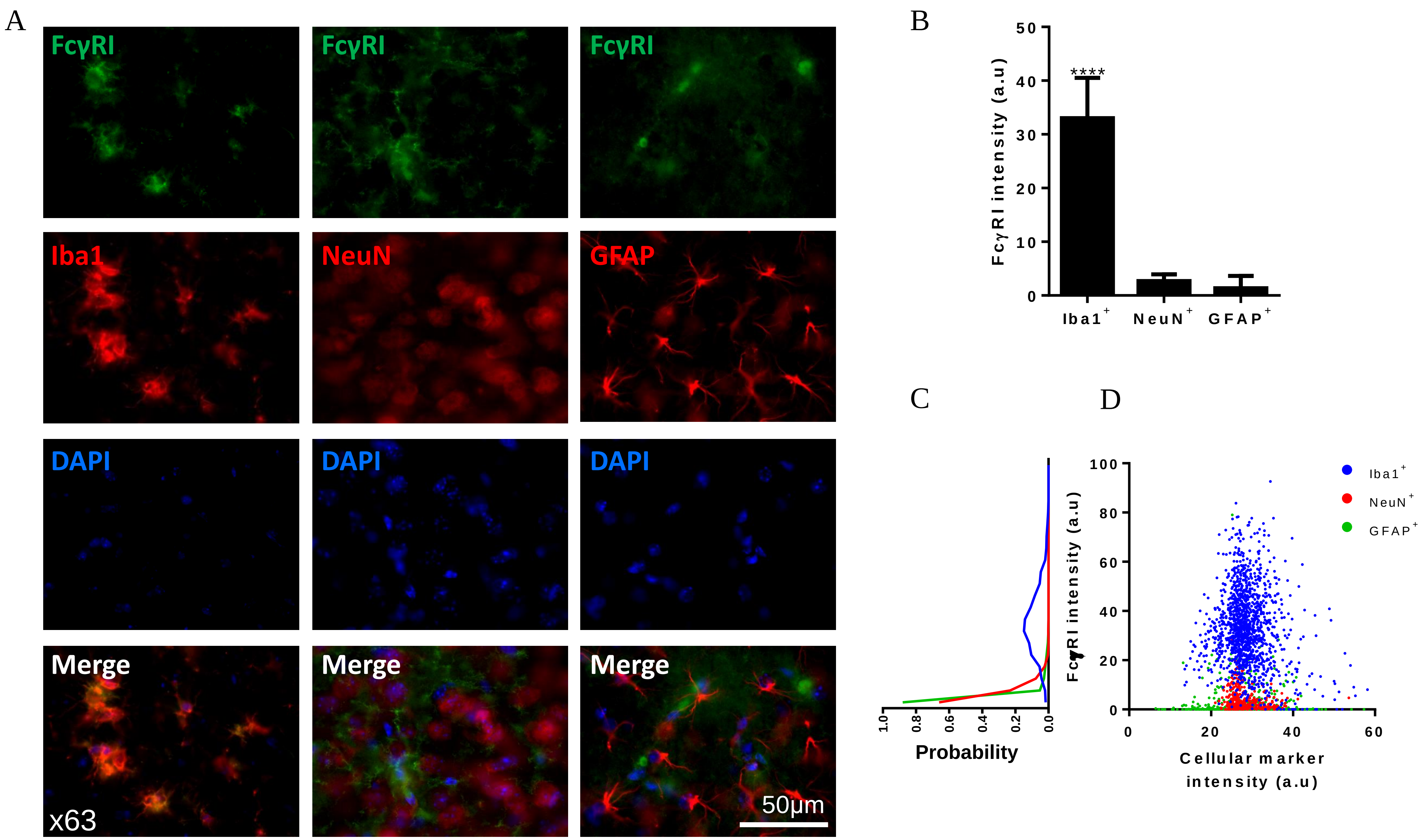
A
FcγRI
FcγRI
FcγRI
Iba1
NeuN
GFAP
DAPI
DAPI
DAPI
Merge
Merge
Merge
x63
50µm
B
FcγRI intensity (a.u)
****
Iba1+
NeuN+
GFAP+
C
Probability
D
FcγRI intensity (a.u)
Cellular marker intensity (a.u)
Iba1+
NeuN+
GFAP+

Figure S7

**Fig. S7. FcγRI is expressed on microglial cells.** To assess FcγRI expression among CNS cells, double labeling was conducted with (A) $Iba1^{+}$ (Microglia), NeuN (mature neurons) and GFAP (astrocytes), using the x10 objective. (B) FcγRI fluorescent signal intensity is higher in $Iba1^{+}$ cells compared with both NeuN and $GFAP^{+}$ cells. (C) Scatter plot and distributions of FcγRI expression on microglia, neurons, and astrocytes, showing normal distribution among microglia and right-skewed distributions for astrocytes and neurons (D) FcγRI is expressed within the somas of Iba1 $^{+}$cells. Images were taken using the x40 (left panels) and x63 (right panels) objectives. ****P<0.0001, one-way ANOVA.

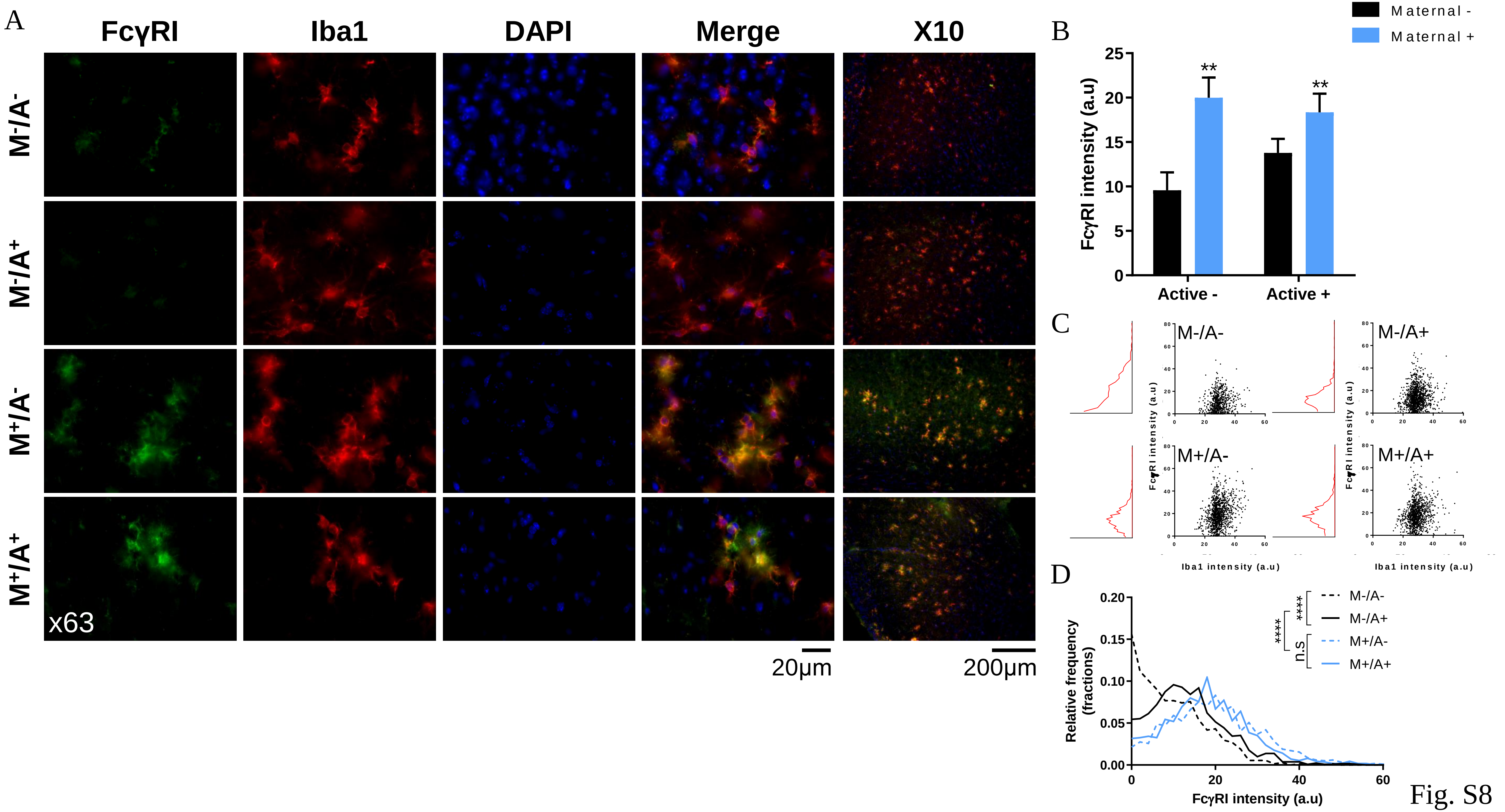
A
FcγRI
Iba1
DAPI
Merge
X10
M-/A-
M-/A+
M+/A-
M+/A+
x63
20μm
200μm
B
Maternal -
Maternal +
FcγRI intensity (a.u)
25
20
15
10
5
0
**
**
Active -
Active +
C
M-/A-
M-/A+
M+/A-
M+/A+
FcγRI intensity (a.u)
Iba1 intensity (a.u)
D
Relative frequency (fractions)
0.20
0.15
0.10
0.05
0.00
0
20
40
60
FcγRI intensity (a.u)
M-/A-
M-/A+
M+/A-
M+/A+
****
****
n.s
Fig. S8

**Fig. S8. FcγRI is upregulated among cortical microglia of maternally vaccinated 5xFAD mice.** (A) FcγRI expression was assessed using double-labeled immunofluorescence with $Iba1^{+}$ microglia, using the x10 and x63 objectives for visualizing and quantification, respectively. (B) FcγRI signal was increased among microglia from maternally vaccinated mice independently of active vaccination. (C) Scatter plot of Iba1 and FcγRI signals reveal right-skewed distribution for FcγRI expression among unvaccinated and actively vaccinated mice, and normal distribution for maternally vaccinated mice. (D) Overlay and comparisons of FcγRI expression distribution. **P<0.01, ****P<0.0001, two-way ANOVA, corrected two-sample Kolmogorov-Smirnov test.

A

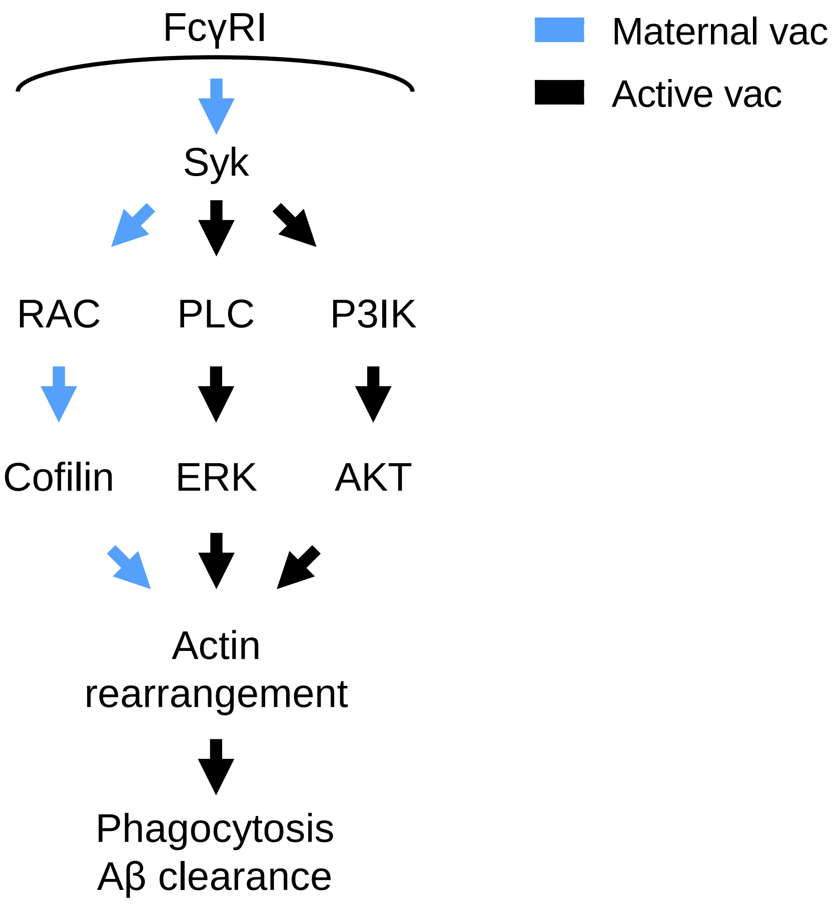

FcγRI
Syk
RAC
PLC
P3IK
Cofilin
ERK
AKT
Actin rearrangement
Phagocytosis
Aβ clearance
Maternal vac
Active vac


Fig. S9

**Fig. S9. Maternal vaccination activates FcR-mediated phagocytosis via activation of AKT, ERK, and Cofilin actin-cytoskeleton regulation pathways.** Levels of Syk and downstream signaling molecules from the Fc-mediated phagocytosis pathway were measured using immunoblotting following maternal vaccination and active vaccination at 5m of age (see main Fig. 5A). (A) Maternal and active vaccination activates FcγR-mediated phagocytosis pathways through actin cytoskeleton regulation.

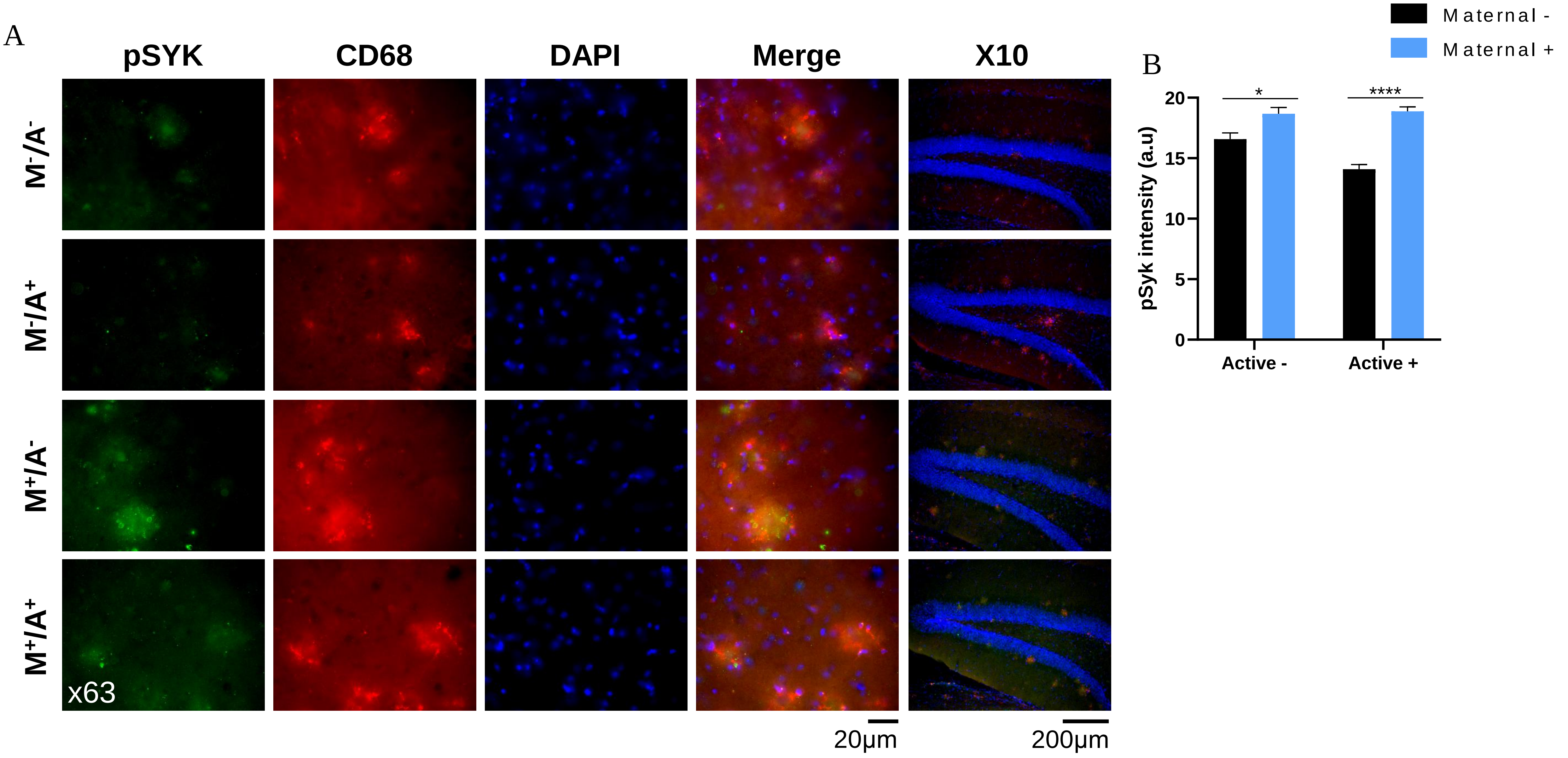
A
pSYK
CD68
DAPI
Merge
X10
M-/A-
M-/A+
M+/A-
M+/A+
x63
20μm
200μm
B
Maternal -
Maternal +
pSyk intensity (a.u)
20
15
10
5
0
*
****
Active -
Active +

Fig. S10

**Fig. S10. Maternal vaccination increases Syk activation in hippocampal microglia.** (A) Syk activation among microglial cells was assessed using double labeling of pSyk and CD68 positive microglia, using the x10 and x63 objectives for visualizing and quantification, respectively. (B) pSyk signal was higher in maternally and actively immunized mice compared with unvaccinated controls. *P<0.05, ****P<0.0001, two-way ANOVA.

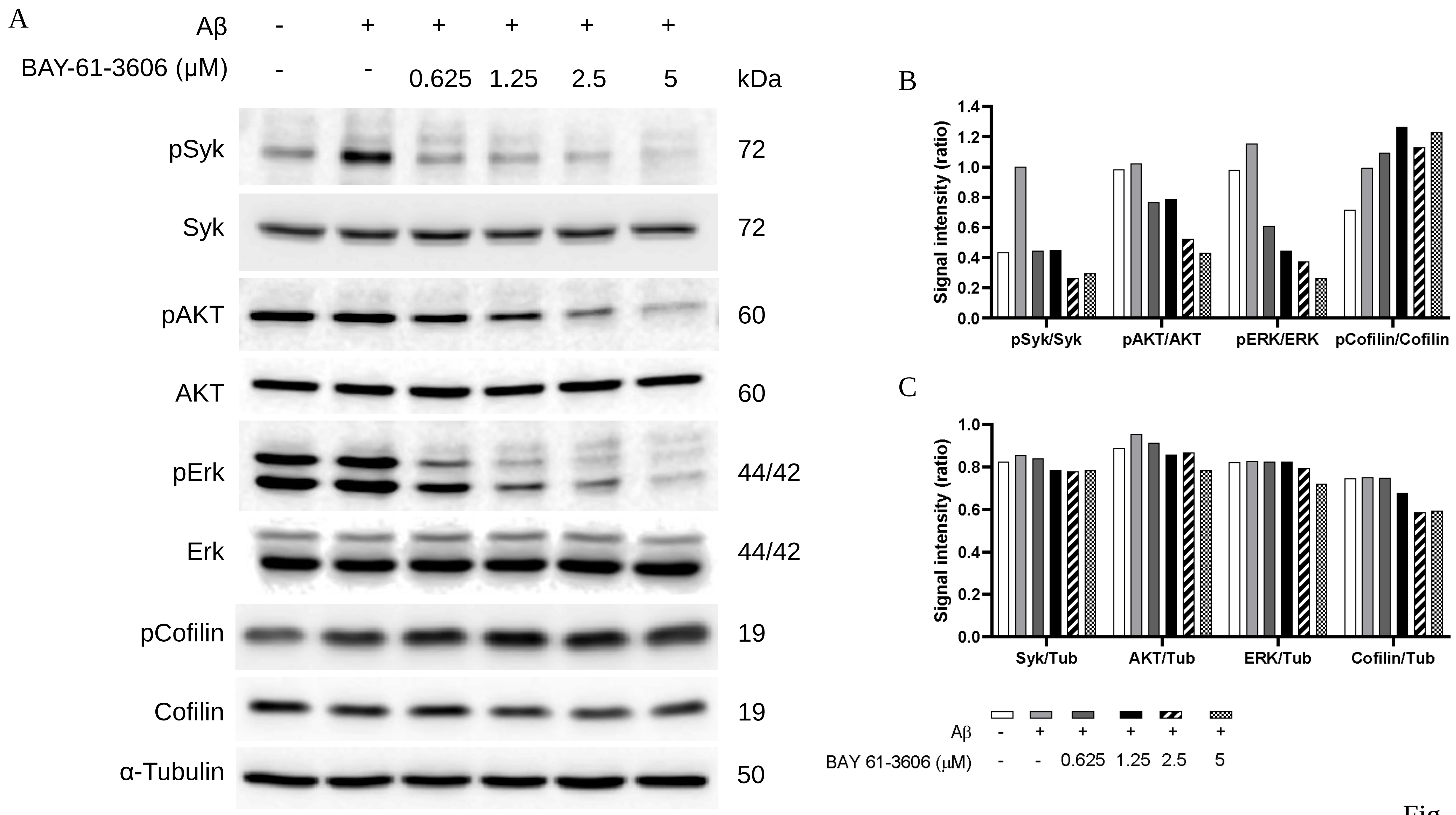
A
Aβ - + + + + +
BAY-61-3606 (μM) - - 0.625 1.25 2.5 5
kDa
pSyk 72
Syk 72
pAKT 60
AKT 60
pErk 44/42
Erk 44/42
pCofilin 19
Cofilin 19
α-Tubulin 50
B
Signal intensity (ratio)
0.0 0.2 0.4 0.6 0.8 1.0 1.2 1.4
pSyk/Syk
pAKT/AKT
pERK/ERK
pCofilin/Cofilin
C
Signal intensity (ratio)
0.0 0.2 0.4 0.6 0.8 1.0
Syk/Tub
AKT/Tub
ERK/Tub
Cofilin/Tub
Aβ - + + + + +
BAY 61-3606 (μM) - - 0.625 1.25 2.5 5

Fig. S11

**Fig. S11. Dose-dependent Syk inhibition by BAY-61-3606 in the N9 microglial cell line.** BAY-61-3606 was applied to N9 cells at different concentrations, ranging from 0.75 to 5μM for 2h, followed by the addition of aggregated human A$\beta_{42}$ peptide at a concentration of 750nM. (A-C) Western blotting of phospho- and total Syk and downstream signaling molecules from the FcR mediated phagocytosis pathway: AKT, ERK, and Cofilin.
