## Supplementary Materials and Methods for "Maternal antibodies facilitate Amyloid-β clearance by activating Fc-receptor-Syk-mediated phagocytosis"

**Vaccine administration.** Mice were injected intramuscularly with 25µg DNA (50µl) and electroporation was administered immediately to the area of the injection using a two-needle array electrode, 10mm (BTX, 45-0167, Holliston, MA) and an ECM830 electroporator (BTX, 45-2052). Electroporation configuration: one pulse of 450V/cm, 2 repetitions, duration=0.05ms, interval=0.125s following a second pulse of 110V/cm, 8 repetitions, duration=10ms, interval=0.125s (*20*).

**Serum collection.** Blood was extracted from the facial vein using a glass cannula and incubated for 30min at RT to clot. Samples were then centrifuged at 1500×g for 8min at 4°C and clear serum was stored at -20°C for further analysis.

**Antibody titer.** Anti-A$\beta_{1-11}$ Ab production was quantified by a standard indirect ELISA. 96-well high binding microplates (Microlon, 655061, Greiner bio-one, Monroe, NC) were covered with 50µl of recombinant mouse A$\beta_{1-11}$ peptide (custom synthesis, Adar-Biotech, Yavne, Israel) in carbonate/bicarbonate coating buffer (pH 9.6) at concentration of 3µg/ml. Plates were incubated overnight at 4°C, washed 3 times with 0.1% PBS-Triton and blocked with 2% Bovine Serum Albumin (BSA) (A7906, Sigma, St. Louis, MO) for 1h at RT. Plates were washed 3 times with PBS-T following serum incubation at dilutions of 1:100-1:12,500 for 1h at RT. Standard curve was carried out using known concentrations of primary rabbit anti-Aβ Aβ1-14 Ab (50-500ng/ml, ab2539, Abcam, Cambridge, UK). Plates were washed 3 time in PBS-T and incubated with HRP-conjugated goat anti-mouse IgG secondary Abs diluted at 1:5,000, (115-035-003, Peroxidase

AffiniPure, Jackson Immunoresearch, PA) or goat anti-rabbit secondary Ab for standard curve wells (111-035-003, Peroxidase AffiniPure, Jackson Immunoresearch) for 1h at RT. Plates were washed 3 times in PBS-T and 3,3',5,5'-tetramethylbenzidine (TMB) substrate (00-4201-56, Affimetrix eBioscience, San Diego, CA) was applied. Colorimetric reaction was stopped by adding 50µl of 2M $H_2SO_4$ solution (339741, Sigma-Aldrich, St. Louis, MO). OD was measured at 450nm using a spectrophotometer.

**Immunoglobulin isotyping.** IgG isotyping was conducted using a similar indirect ELISA protocol with the addition of specific anti-mouse-immunoglobulin Abs (ISO-2, Sigma-Aldrich, St. Louis, MO) diluted at 1:1000, incubated for 30min at RT. Next, donkey anti-goat secondary Ab (705-035-003, Peroxidase AffiniPure Donkey Anti-Goat IgG, Jackson Immunoresearch) diluted at 1:5,000 was applied for 1h at RT.

**Spatial short-term memory.** We utilized a variant of the T-maze alternation test modified from (*56*). Briefly, T-maze arms were 30cm long and 15cm wide, walls were 15cm high, covered by different black and white patterns. Mice were given 3 trials with a 2hrs inter-trial interval. Each trial consisted of 2 stages: During acquisition, mice were released from the starting chamber and were given the opportunity to enter one of the target arms. A trial was ended when the animals spent more than 2s with all 4 limbs inside one of the target arms. Next, mice were allowed to stay in the chosen arm for 30s, followed by a repetitive trial in which alternation rate was measured.

**Measuring A$\beta_{40/42}$ levels using sELISA.** A$\beta_{40}$ and A$\beta_{42}$ in the cortex were measured using a modification of a previously published sandwich-ELISA protocol (*32*). Briefly, tissues were mechanically homogenized in in TBS-Triton 1% (120mM NaCl, 50mM Tris, pH=8.0, 150mg/ml (tissue/buffer) including protease inhibitor cocktail (1:100, P2714, Sigma, St. Louis, MO), then incubated on ice for 30min followed by centrifugation for 120min at 17,000g at 4 °C. Supernatant, containing TBS-T-soluble fraction of $A\beta_{40}$ and $A\beta_{42}$ was removed and stored at−20°C. The centrifuged pellet was incubated for 30min with 2% TBS-SDS (120mM NaCl, 50mM Tris + 2% SDS) on ice, following centrifugation at the same conditions as mentioned above. Supernatant, containing the SDS-soluble fraction of Aβ was removed and stored at −20°C. The remaining pellet, containing insoluble Aβ was resuspended in 70% formic acid and incubated on ice for 30m, followed by centrifugation at the same conditions as mentioned above. Formic acid-soluble supernatant was separated and neutralized using 1M Tris (pH=11, 20-time the volume of the formic acid) and stored at −20°C. Total protein concentration was determined using the BCA method (Cat#23225, Thermo Fisher Scientific, Waltham, MA). For the ELISA assay, 96-well polystyrene microplates (655061, Greinerbio-one, Monroe, NC) were covered with 50µl of anti-rabbit-N-terminus $A\beta_{1\text{-}14}$ (ab2539, Abcam, Cambridge, UK) at a concentration of 5µg/ml in carbonate-bicarbonate buffer (pH=9.6) and incubated overnight at 4°C. Plates were washed 4 times in PBS-T solution (0.1% Triton-x in PBS) and blocked with 2% BSA solution in PBS. 50µl of tissue homogenate were applied to each well, and incubated for 60min at RT. Plates were then washed 5 times in PBS-T, and the following detection Abs were added: Anti-$A\beta_{40}$ Ab (ab20068, Abcam, Cambridge, UK) diluted at 1:500 or anti $A\beta_{42}$ Ab (05-831, Millipore, Billerica, MA) at 1:2500, and incubated for 60 min at RT. Next, plates were washed 5 times in PBS-T and secondary goat-anti-mouse IgG HRP-conjugated Ab was added (115-035-003, Peroxidase AffiniPure Goat Anti-Mouse, Jackson immunoresearch, PA) at a dilution of 1:5,000. Plates were washed 5 times with PBS-T and 50µl of 3, 3', 5, 5'-tetramethylbenzidine (TMB) substrate (00-4201-56, Affimetrix eBioscience, San Diego, CA) was added. The color reaction was allowed to develop for 3 min and was stopped by adding 50µl of 2M H2SO4. Optical density (OD) was measured at 450nm using a spectrophotometer. Standard curve was carried out using known concentrations of recombinant $A\beta_{40}$ and $A\beta_{42}$.

**RT-qPCR.** Total RNA was extracted using TRIzol Reagent (Ambion, Life Technologies, CA). Complementary DNA (cDNA) was generated using Revert Aid H minus first strand cDNA synthesis kit (Thermo Scientific, Waltham, MA). RT-PCR reactions were performed using Fast SYBR Green Master Mix (Applied Biosystems, CA) in a StepOnePlus instrument (Applied Biosystems, CA). Primers (Supplementary table 2) were calibrated, a negative control was performed for each primer pair and PCR products were validated in gel electrophoresis. Samples were measured in triplicates and values were normalized according to mRNA levels of β-Actin. Denaturation was performed at 94°C for 30s, annealing for 10s and elongation was performed at 72°C for 10s.

**Western blot.** Hippocampal protein lysate was obtained as mentioned above with the addition of phosphatase inhibitor cocktail (1:100, 524625, Merck-Millipore, Billerica, MA). 25µg per of total protein per sample was boiled in Laemmli buffer at 95°C for 10min and loaded to 10% (w/v) Tris-glycine polyacrylamide gels. Electrophoresed samples were transferred to a PDVF (IPVH00010 Immobilon-P Membrane, PVDF, 0.45µm, Merck, Kenilworth, NJ) and blocked for unspecific binding using 5% BSA (A7906, Sigma, St. Louis, MO) for phospho-proteins and 5% skim milk (M530, Himedia, Mumbai, India) for unphosphorylated proteins, diluted in 0.1% (v/v) PBS-Tween20 (0.1% Tween 20, P9416-50ML, Sigma, St. Louis, MO). Next, membranes were incubated with a primary Ab overnight at 4°C. Full Abs details can be found in Supplementary table 1. Unbound Abs were washed 3 times in PBS-T for 5min followed by membrane incubation with HRP-conjugated goat anti mouse IgG secondary Ab (Cat#115-035-003, Peroxidase AffiniPure, Jackson immunoresearch) or goat anti-rabbit secondary Ab (111-035-003, Peroxidase AffiniPure, Jackson Immunoresearch) diluted at 1:10,000 in blocking buffer for 1h at room temperature (RT). Ab-Ag bindings were detected by applying ECL (ECL kit, 20-500-120, Biological industries, Israel).
