## Supplementary Tables for "Maternal antibodies facilitate Amyloid-β clearance by activating Fc-receptor-Syk-mediated phagocytosis"

**Supplementary Table 1. sELISA, IF and WB primary antibodies**

| *Protein* | *Application* | *Catalog #* | *Manufacturer* | *Dilution* | *Host* | *Secondary Ab* | *Manufacturer* | *Dilution* |
|---|---|---|---|---|---|---|---|---|
| $A\beta_{1\text{-}14}$ | sELISA (capture) | ab2539 | Abcam, Cambridge, UK | 1:200 | Rabbit | N/A | | |
| $A\beta_{1\text{-}40}$ | sELISA (detection) | ab20068 | Abcam, Cambridge, UK | 1:500 | Mouse | Anti-mouse IgG, HRP | 115-035-003, Jackson immunoresearch | 1:5000 |
| $A\beta_{1\text{-}42}$ | sELISA (detection) | 05-381-I | Millipore, Billerica, MA | 1:2000 | Mouse | Anti-mouse IgG, HRP | 115-035-003, Jackson immunoresearch | 1:5000 |
| $A\beta_{1\text{-}42}$ | IF | 05-381-I | Millipore, Billerica, MA | 1:1000 | Mouse | Anti-mouse IgG, Alexa-488/568 | Invitrogen | 1:1000 |
| Iba1 | IF | 019-19741 | Wako, Osaka, Japan | 1:1000 | Rabbit | Anti-rabbit IgG, Alexa-488/568/647 | Invitrogen | 1:1000 |
| NeuN | IF | MAB377 | Millipore, Billerica, MA | 1:10000 | Mouse | Anti-mouse IgG, Alexa-488/568 | Invitrogen | 1:1000 |
| GFAP | IF | M0761 | Agilent, Santa-Clara, CA | 1:7500 | Rabbit | Anti-rabbit IgG, Alexa-488/568/647 | Invitrogen | 1:1000 |
| FcγRI | IF | MCA5997 | Bio-Rad, Hercules, CA | 1:1000 | Rat | Anti-rat IgG, Alexa-488/568 | Invitrogen | 1:1000 |
| FcγRIIb | IF | MCA6001 | Bio-Rad, Hercules, CA | 1:200 | Rat | Anti-rat IgG, Alexa-488/568 | Invitrogen | 1:1000 |

| | | | | | | | | |
|---|---|---|---|---|---|---|---|---|
| FcγRIII | IF | MCA5998 | Bio-Rad, Hercules, CA | 1:1000 | Rat | Anti-rat IgG, Alexa-488/568 | Invitrogen | 1:1000 |
| FcγRIV | IF | MCA5999 | Bio-Rad, Hercules, CA | 1:1000 | Rat | Anti-rat IgG, Alexa-488/568 | Invitrogen | 1:1000 |
| CD68 | IF | ab53444 | Abcam, Cambridge, UK | 1:2250 | Rat | Anti-rat IgG, Alexa-488/568 | Invitrogen | 1:1000 |
| pSyk | IF | CST-2710 | Cell-Signaling, Danvers, MA | 1:500 | Rabbit | Anti-rabbit IgG, Alexa-488/568/647 | Invitrogen | 1:1000 |
| pSyk | WB | CST-2710 | Cell-Signaling, Danvers, MA | 1:1000 | Rabbit | Anti-rabbit IgG, HRP | 111-035-003, Jackson Immunoresearch | 1:10000 |
| Syk | WB | CST-13198 | Cell-Signaling, Danvers, MA | 1:1000 | Rabbit | Anti-rabbit IgG, HRP | 111-035-003, Jackson Immunoresearch | 1:10000 |
| pAKT | WB | CST-4060 | Cell-Signaling, Danvers, MA | 1:2000 | Rabbit | Anti-rabbit IgG, HRP | 111-035-003, Jackson Immunoresearch | 1:10000 |
| AKT | WB | CST-2920 | Cell-Signaling, Danvers, MA | 1:2000 | Mouse | Anti-mouse IgG, HRP | 115-035-003, Jackson immunoresearch | 1:10000 |
| pERK | WB | CST-4370 | Cell-Signaling, Danvers, MA | 1:2000 | Rabbit | Anti-rabbit IgG, HRP | 111-035-003, Jackson Immunoresearch | 1:10000 |
| ERK | WB | CST-4696 | Cell-Signaling, Danvers, MA | 1:2000 | Mouse | Anti-mouse IgG, HRP | 115-035-003, Jackson immunoresearch | 1:10000 |
| β-Tubulin | WB | T5076 | Sigma-Aldrich, St. Louis, MO | 1:10000 | Mouse | Anti-mouse IgG, HRP | 115-035-003, Jackson immunoresearch | 1:10000 |

| | | | | | | | | |
|---|---|---|---|---|---|---|---|---|
| pCofilin | WB | CST-3313 | Cell-Signaling, Danvers, MA | 1:1000 | Rabbit | Anti-rabbit IgG, HRP | 111-035-003, Jackson Immunoresearch | 1:10000 |
| Cofilin | WB | ab54532 | Abcam, Cambridge, UK | 1:500 | Mouse | Anti-mouse IgG, HRP | 115-035-003, Jackson immunoresearch | 1:10000 |
| β-Actin | WB | sc-47778 | Santa-Cruz Biotechnology, Dallas, TX | 1:1000 | Mouse | Anti-mouse IgG, HRP | 115-035-003, Jackson immunoresearch | 1:10000 |
| hAPP | WB | 803001 | Biolegend, San Diego, CA | 1:5,000 | Mouse | Anti-mouse IgG, HRP | 115-035-003, Jackson immunoresearch | 1:10000 |

**Supplementary Table 2. RT-PCR primers**

| *Gene* | *Forward primer* | *Reverse primer* | *Annealing temp (°C)* | *Product size (bp)* |
|---|---|---|---|---|
| *FCGR1* | AGGTTCCTCAATGCCAAGTG | ATTCTTCCATCCGTGACACC | 60 | 127 |
| *FCGR3* | TATCGGTGTCAAATGGAGCA | TATGGCACCTTAGCGTGATG | 60 | 130 |
| *FCGR4* | CGAGGACAATTCTATCAAGTGGTT | ACTTAGTGGTCTGAAGCAATAGCC | 60 | 188 |
| *FCRN* | ACTGCTAGGCCACCTGGAG | AGGAGAAAGCAGCACAGGTC | 64 | 122 |
| *CD68* | ACTTCGGGCCATGTTTCTCT | GCTGGTAGGTTGATTGTCGT | 60 | 138 |
| *ACTIN* | TTCTTTGCAGCTCCTTCGTT | ATGGAGGGGAATACAGCCC | 56 | 149 |
